# Stability of neural manifolds at minimal dimensionality despite motor representational drift

**DOI:** 10.64898/2026.01.30.702162

**Authors:** Muhammad U. Abdulla, Stephen E. Clarke, Elizabeth J. Jun, Paul Nuyujukian Neural Prosthetic Systems Lab and the Brain Interfacing Lab

## Abstract

Recent work has modeled the generation of movement as emerging from a latent dynamical system. While the relationship between recorded neural populations and latent variables is non-stationary, latent trajectories populate low-dimensional manifolds that appear stable over time. However, the dimensionality of these manifolds and their relationship to motor cortical circuitry remains unclear. We propose a simple framework for extracting labeled latent variables that maintain a fixed relationship to movement parameters in a two-dimensional reaching task. Despite only capturing 3-7% of total variance and spanning two dimensions, this supervised method outperforms other common methods at an offline decoding task, and explains the long-term stability of neural manifolds observed in previous literature. We demonstrate that changes in the *encoding map* (from neurons to latents) is the dominant source of drift in motor cortex, while the *latent map* (from latents to behavior) remains stable over years. Additionally, we find that, for a two-dimensional task, neural structure is constrained by task complexity, limiting the insights that neural manifolds offer onto underlying circuitry. Our results motivate future studies to uncover causal relationships between neural computation and behavior.

## Introduction

A primary goal of systems neuroscience is to model the relationship between neural activity and behavior. Initial work in the field sought to identify direct correlations between stimuli and the spiking activity of individual neurons ^1–5^. Nonetheless, the characterization of neuronal tuning remained elusive in several regions of the brain, such as CA2, where neural activity patterns change over time ^6^, and M1, where correlations to behavior are dynamic over the duration of a motor task ^7^. This characterization is further complicated by non-stationary correlations between neural activity and experimental conditions, referred to as *representational drift*, which have been observed in both deep brain ^6,8,9^ and cortical regions ^10–15^. It has been hypothesized that drift is a general phenomenon allowing the brain to continually learn ^12–14^ and maintain a balance between plasticity and stability ^16^.

Over the past two decades, the dominant paradigm in motor systems neuroscience has shifted from emphasizing how individual neurons encode specific motor functions ^4,17^ to how populations of neurons generate movement ^18–21^. Early work demonstrated that neural populations align with preferred activity patterns during the preparatory phase of voluntary reaches ^22,23^. The subsequent neural state dynamics are highly stereotyped with respect to conditions of experimental tasks ^22–24^, involve oscillatory components ^25^, and can be modeled accurately using recurrent neural network architectures ^19,26,27^. While the circuitry involved in these dynamics spans millions of neurons, observing just a small number of motor units is often sufficient for characterizing activity patterns hypothesized to generate movement ^18^. Recent literature has advanced this perspective by developing new methods for extracting latent variables from recorded neural activity ^28–33^ and modeling their relationship to motor outputs ^28,34–36^.

In the context of brain-machine interfaces (BMIs), simple linear methods have been sufficient for inferring motor parameters from experimental tasks in real-time (e.g. hand position during a center-out reaching task) ^37,38^. For instance, projecting neural activity onto the linear subspace spanned by the 10 dominant eigenvectors identified by principal component analysis (PCA) results in latent trajectories that populate a low-dimensional manifold and maintain correlations to hand velocity during voluntary reaches ^39^. Moreover, these latent trajectories appear stable over time, as supervised methods such as canonical correlation analysis (CCA) can be used to align latent embeddings between experimental sessions ^39^. In other systems, stable neural manifolds have served as powerful abstractions that offer insights onto the underlying circuitry ^40^ (e.g. the head-direction system ^41^). Thus, it has been hypothesized that the stability of approximately 10-dimensional neural manifolds (with estimates of dimensionality ranging from 5–20 ^7,25,28,36,39,42,43^, see Supplementary Table 1) in motor regions arises from constraints of the neural circuitry that generates movement ^36,39,42,44–46^.

In this work, we propose a simple framework for characterizing the relationship between recorded neural population activity and motor execution as the composition of an *encoding map* that transforms neural population activity to a latent space, and a *latent map* that transforms latent variables to movement parameters. This approach allowed us to separate potential sources of stability and drift. By applying a supervised dimensionality reduction method, we consistently isolated a latent space with a fixed relationship to behavior in the minimum number of dimensions allowed by task-constraints ^47,48^. We demonstrated that a two-dimensional latent space was sufficient to perform stable offline decoding of movement parameters over years of experiments, a significantly lower estimate for the dimensionality of the behaviorally relevant latent space than previously reported. Using this framework, we propose that preserved latent stability might arise as a byproduct of regressing a large, drifting population of neurons against relatively simple and consistent tasks with constrained topology, which potentially limits the insights offered onto the topology of the underlying circuitry. We argue that prior frameworks struggle to sufficiently distinguish these two explanations of latent stability, suggesting a reevaluation of experimental methodology and the respective mathematical modeling underlying the neural control of movement.

## Results

### Model hypothesis

Data was collected from four male rhesus macaques performing an eight-target center-out reaching task on a computer monitor, with spiking data in motor cortex (M1) recorded over *n*_ch_ = 96 channels on a microelectrode array (for further details, see Methods). As depicted in Figure 1, the proposed framework consists of:

**Figure 1:**
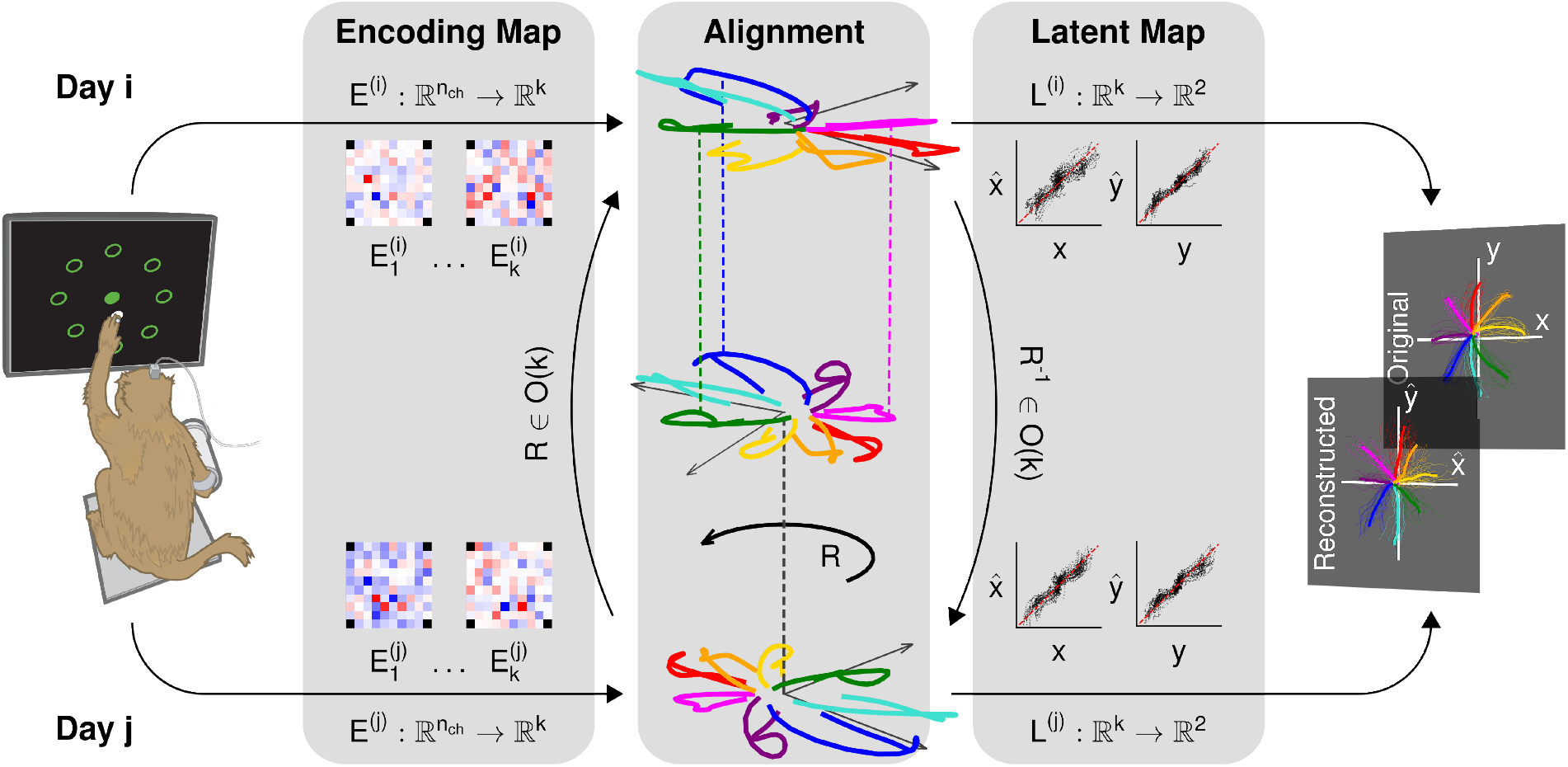
A proposed framework for mapping neural activity to behavior. Rhesus macaques performed an eight-target, center-out reaching task on a computer monitor while neural and kinematic activity was recorded concurrently. For each experimental session, we identified an *encoding map E*, that projected population activity onto a *k*-dimensional latent space, and a *latent map L*, that transformed the latent space to behavioral outputs. Square arrays in the “Encoding Map” block indicate the relative weight that encoding maps apply to each electrode on the array, which changes between Day *i* and Day *j*. Trajectories in the “Alignment” block depict the average latent trajectory for each reach direction in the 2-dimensional subspace identified using multiclass linear discriminant analysis (LDA) for both Days *i* and *j*. A low-dimensional rotation *R* ∈ *O*(*k*) was sufficient to align latent trajectories between experimental sessions. Thus, the latent map trained on Day *i* could be applied to latent dynamics extracted from Day *j* to reconstruct cursor kinematics. Days *i* and *j* in this figure correspond to datasets J110902 01 and J130620 01, respectively, recorded 1.8 years apart. Scatter plots depicted in the “Latent Map” block show results of least squares regression used to train a velocity-based linear control system to reproduce cursor positions from recorded neural data.

1. An *encoding map E* that performs a linear transformation from population activity recorded across *n*_ch_ channels to a *k*-dimensional latent space. For experimental data recorded on Day *i*, the encoding map is identified as an *n*_ch_-by-*k* matrix, *E*^(*i*)^. The columns 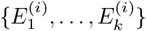 of the encoding map are patterns of covariance, or *encoding patterns*, of neural spiking data from the array. The dynamical systems view models movement as being generated by flexible combinations of populations of neurons in motor regions ^18^. As in past literature, relevant states of population activity are approximated using linear combinations of activity across the microelectrode array. The encoding map projects a neural population state *n*(*t*) to a latent state *ℓ*(*t*) = *E*^(*i*)*T*^ *n*(*t*), where the dominant patterns of neural activity at time *t* have the largest values in the resulting latent state vector. Additionally, we constrain the columns of the encoding map to be orthonormal (a common constraint for linear dimensionality reduction methods, see Methods), guaranteeing that patterns are linearly independent and relative weights of latent variables reflect the relative prominence of corresponding encoding patterns.
2. A *latent map*, which transforms the latent state to a desired output. A latent map is generated for each of the following tasks:
  - a *decoding* task, where the goal is to infer the task condition (i.e. the intended reach direction) from samples in the latent space. In this case, the latent map is a classifier trained on the conditional probability distribution of the latent state with respect to each reach direction, approximated as a Gaussian.
  - a *control* task, where the true cursor kinematics are reconstructed using a control system. In this case, the latent map is an affine function *L* : ℝ^*k*^→ ℝ^2^ transforming the latent state into an instantaneous cursor velocity vector, which can be integrated with respect to time to reconstruct cursor position (see Methods for more details).

This proposed framework builds on previous motor systems neuroscience literature. Identifying an encoding map is equivalent to the well-established goal of extracting relevant latent variables from recorded population activity. Gradual changes of this encoding map should be expected, due to changes in how the recorded subpopulation of cortical neurons correlate to the underlying dynamical state (i.e. representational drift ^12,14,15^) and sampling instability of the recording device ^39,49^. Similarly, training a latent map is equivalent to characterizing a precise relationship between latent variables and movement parameters (a frequent objective in BMI literature ^26,37,38^). In past work, a *k* = 10-dimensional linear subspace was identified using PCA, which resulted in latent trajectories that correlated to hand velocities. Furthermore, a supervised approach (CCA) was used to align latent variables between experimental sessions, maintaining kinematic decoding performance over time ^39^. It has been hypothesized that the stability of latent dynamics and correlations to behavior in the identified linear subspace emerge (at least in part) from constraints imposed by neural circuitry, and therefore that properties of this *intrinsic manifold* effectively capture the structure of underlying circuitry ^39,42,44,46^. However, there are many alternative candidate explanations for this observed stability; for instance, latent trajectories between experimental sessions could be highly correlated to one another due to shared correlations to kinematic behavior while performing the same task. More formally, let *L* : ℝ^*k*^ → ℝ^2^ be a linear regression map that approximates hand velocities from latent states, and assume that these quantities are sufficiently correlated. Using QR-decomposition, this map can be decomposed as *L* = *Q*_*L*_*R*_*L*_, where *Q*_*L*_ projects latent variables to a 2-dimensional subspace, and *R*_*L*_ is a homeomorphism between this “regression subspace” and hand velocities. Since topologies are preserved under homeomorphisms, latent trajectories in the regression subspace would have a similar topology to hand velocities. Moreover, hand velocity traces are nearly identical between experimental sessions since the same motor task is being performed. Latent trajectories in the regression subspaces from different experimental sessions would then share similar topology to corresponding hand velocities, and thus to each other. If the 10-dimensional PCA subspace consistently spans the 2-dimensional regression subspace, then it is reasonable to expect that supervised methods (such as CCA) could accurately align regression subspaces between sessions, allowing for consistent decoding of behavior. Importantly, this would result in a stable relationship between neural manifolds and motor outputs solely due to correlations and constraints imposed by the experimental task, not necessarily the underlying neural circuitry.

To evaluate how latent stability and offline BMI performance changes as a function of latent space dimensionality, we propose using multiclass linear discriminant analysis (LDA) ^50^ to identify an encoding map of dimension *k* ∈ { 1, …, 7 }. This results in a subspace where reaches in different directions are separated, while reaches in the same direction follow similar trajectories. These properties are relevant to the encoding of motor control, since underlying neural states are known to align with preferred states during the preparatory phase of reaching, and subsequently follow stereotyped dynamics ^23,24^. Furthermore, unlike linear regression, the dimensionality of the resulting subspace is constrained by the number of task conditions (i.e. choices for reach directions), not the degrees of freedom of the recorded movement (see Methods). In the following sections, we verify that (within the context of 96-channel microelectrode array recordings in M1) this LDA-subspace contains sufficient information for classifying task conditions and reproducing motor kinematics. We demonstrate that projecting to a latent space of only 2-dimensions is indeed sufficient for building an offline BMI with a fixed latent map, which achieves stable performance over months and years across multiple subjects. We compare the performance of the method proposed in this work to past work on identifying stable latent representations ^39^, particularly with respect to the assumed dimensionality of the intrinsic manifold. Finally, we propose a method for quantifying representational drift, and demonstrate that drift of the encoding map is the dominant source of non-stationarity in M1.

### LDA identifies a behaviorally relevant latent space with minimal dimensionality

We compared four linear dimensionality reduction methods to determine which identified the best encoding map: (1) multiclass linear discriminant analysis (LDA) with orthogonalization; (2) principal component analysis (PCA), which identifies dominant sources of total variance; (3) demixed principal component analysis (dPCA), which identifies dominant sources of condition-dependent variance ^51^; and (4) canonical correlation analysis (CCA), which identifies a subspace that is highly correlated to kinematic data. For simplicity, we limited our consideration to linear methods, since previous literature has established they are often sufficient for identifying neural manifolds ^39,42,44,45^ and training BMIs ^37,38^. In Figure 2, we depict the performance of each method across multiple criteria: (1) an offline decoder task, where reach direction was inferred from instantaneous latent states using a Gaussian Naive-Bayes decoder; (2) the proportion of total neural spiking variance explained by the identified subspace; (3) and a control task, where observed cursor kinematics were reconstructed using a linear control system. A 5 × 2-cross validation ^52^ was used to estimate the distribution of decoder accuracy rates, proportions of variance explained, and coefficients of determination between true and reconstructed kinematics for each of four Monkeys (*J, U, L*, and *O*), across all linear methods and subspace dimensionalities. Performance using alternative evaluation metrics is depicted in Supplementary Figure 2.

**Figure 2:**
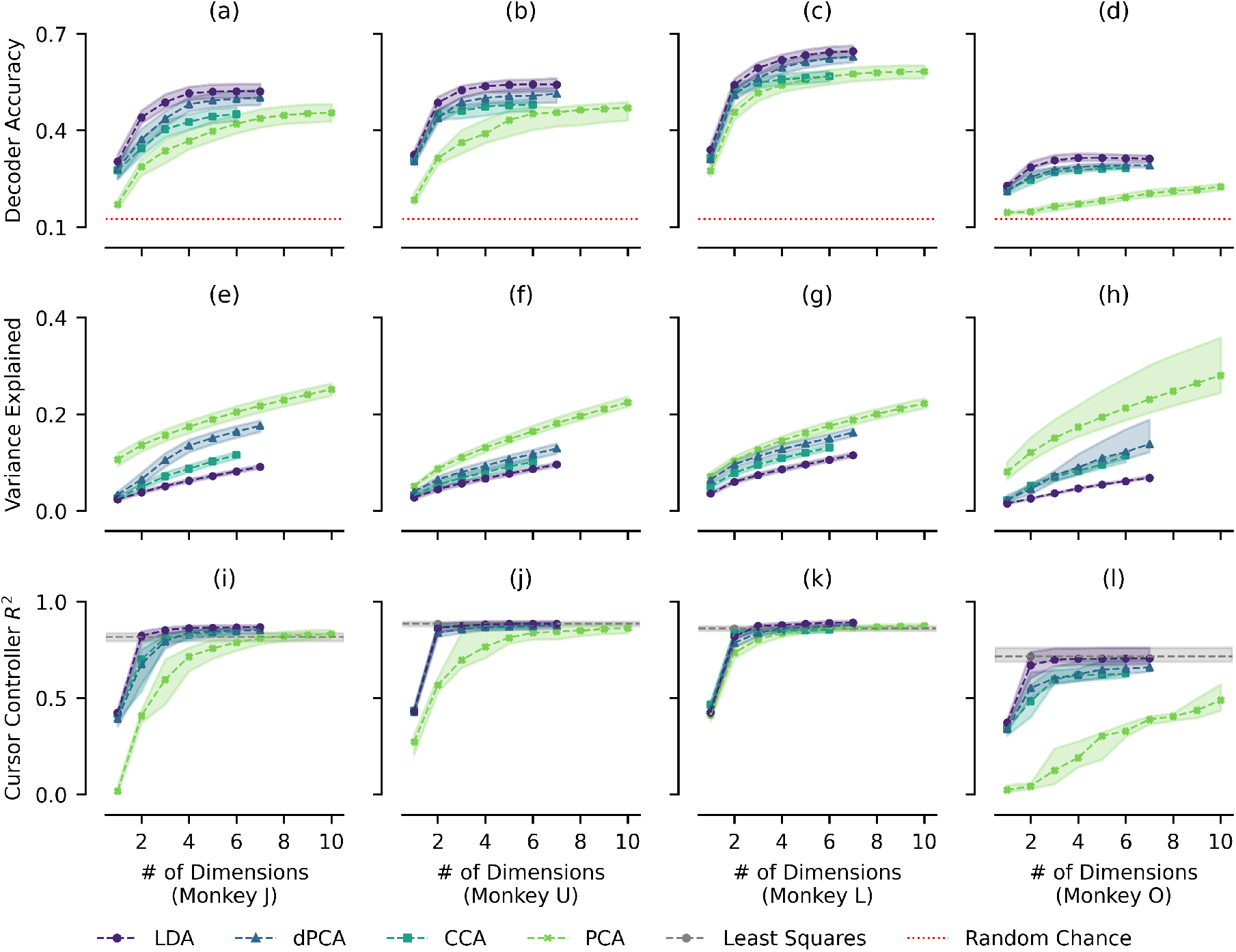
Performance of dimensionality reduction methods at identifying behaviorally relevant latent variables vs. subspace dimensionality. Columns indicate results for Monkeys *J* (a, e, i), *U* (b, f, j), *L* (c, g, k), and *O* (d, h, l). Rows indicate how well identified latent variables performed with respect to accuracy of a Gaussian Naive-Bayes decoder (a, b, c, d), proportion of total variance explained (e, f, g, h), and coefficient of determination between true and reconstructed kinematics (i, j, k, l). Performance was evaluated for latent variables identified using LDA (circle, purple), dPCA (triangle, blue), CCA (square, teal), and PCA (X, light green). Points and dotted-lines represent median performance for each subspace dimensionality; shaded regions represent the interquartile range. Row 1 includes random chance performance for decoding reach directions (dotted-line, red). Row 3 includes performance for least squares controller trained on the entire neural population rather than a subspace (dotted-line, gray). Data was taken from all experimental sessions described in Supplementary Table 2. We find that across all monkeys, LDA consistently identified the latent space that performed better at decoder and controller tasks, in a lower number of dimensions, while explaining significantly less total neural spiking variance.

The latent variables identified by LDA (circle, purple) consistently out-performed other methods in terms of decoder accuracy (Figure 2, Row 1) and reconstruction of cursor kinematics (Figure 2, Row 3). This trend persisted across all monkeys and subspace dimensionalities. Furthermore, LDA decoder performance plateaued at 3–4 dimensions for the decoder task, and only 2 dimensions for the controller task. In previous literature, relevant dynamics of motor regions were modeled as occupying a manifold spanning roughly 10-dimensions ^25,28,36,39,42,43^, which was theorized to emerge from constraints of the underlying circuitry ^19,36,39,42,44–46^. It was also assumed that the subspace spanned by the ten largest principal components was a sufficient approximation of this neural manifold ^39^. However, our results indicate that the manifold containing the relevant information for decoding and cursor control in a two-dimensional reaching task spans substantially fewer dimensions and lower proportion of total variance (Figure 2, Row 2) than previously thought. Across all considered linear methods, increasing subspace dimensionality did not vastly improve performance when compared to the 2-dimensional subspace identified via LDA. Distributions of differences in performances across different subspace dimensionalities are depicted in Supplementary Figures 3, 4, 5, & 6.

The performance of dPCA (triangle, blue) converged to the performance of LDA as the subspace dimensionality increased. This is likely because dPCA maximizes the proportion of condition-dependent variance, which biases it towards dominant sources of total variance, i.e. larger principal components. This bias is further evidenced by the fact that dPCA consistently had the second greatest proportion of total variance explained (Figure 2, Row 2), after only PCA (which achieves the upper bound.) Meanwhile, LDA consistently achieved better performance despite capturing significantly less total variance. As discussed in the Methods, discriminant analysis is deeply related to dPCA, and can be approximated as a generalized version of dPCA with an additional regularization term that removes bias towards dominant sources of total variance. Thus, LDA accounts for less total variance explained than other linear methods, and effectively isolates the behaviorally relevant patterns of activity that directly contribute to discrimination between movement parameters.

In the control task (Figure 2, Row 3), projecting the data to a 2-dimensional space using LDA resulted in a controller with comparable (and sometimes better) performance than the least squares controller (dotted, grey) trained on all channels of neural data. This implies that projecting onto the LDA subspace does not discard information necessary to train a linear control system, and might actually prevent over-training on irrelevant activity patterns present in the training set. While many other methods for identifying latent vectors exist, LDA is a simple linear framework that consistently isolates a subspace relevant to the encoding of motor control in a relatively low number of dimensions.

### Long-term stability of the latent space

Given pairs of experimental sessions on Days *i* ≠ *j*, the decoder and control systems trained on Day *i* are evaluated on data from Day *j* under three hypotheses regarding stability and drift in M1.

1. No drift occurs: the encoding and latent maps both remain fixed. This is equivalent to modeling a static relationship between recorded neural activity and behavior.
2. Latent drift: the encoding map remains stable, but the latent map (i.e. the relationship between latent states and behavior) changes over time. Data from both Days *i* and *j* are split into both training and test sets. Data from Day *j* is projected onto the LDA subspace derived from the training set on Day *i*. New latent maps (a Gaussian Naive Bayes decoder and velocity-based control system) are trained on projected training data from Day *j* and evaluated on projected test data from Day *j*. Note that this does not require additional alignment of the latent space, since re-training the latent map accounts for changes in latent trajectories.
3. Encoding drift: the encoding map (i.e. the behaviorally relevant subspace) changes over time, but the latent map remains stable. Data from the training sets on both Days *i* and *j* are projected onto corresponding LDA subspaces. The optimal low-dimensional rotation matrix *R* ∈ *O*(*k*) that best aligns the average trajectories for each reach direction from the training set on Day *j* to those from the training set on Day *i* is identified. (This differs from past literature where general linear transformations were identified using CCA^39^). Projected data from the test set on Day *j* is rotated according to the alignment rotation *R*. The performance of latent maps from the training set on Day *i* are evaluated on aligned data from the test set on Day *j*. Note that constraining alignment maps to rotation matrices does not alter the subspace identified by the encoding map or the resulting “power” of the latent vectors that population activity is projected onto.

Encoding maps were identified using LDA with dimensionality *k* = 2, as this is where performance plateaued in the previous section, and matches the theorized lower bound on dimensionality imposed by constraints of the two-dimensional motor task ^47,48^. Performance at decoder and controller tasks are depicted in Figures 3 & 4, respectively. Additional evaluation metrics are included in Supplementary Figures 7 & 8.

**Figure 3:**
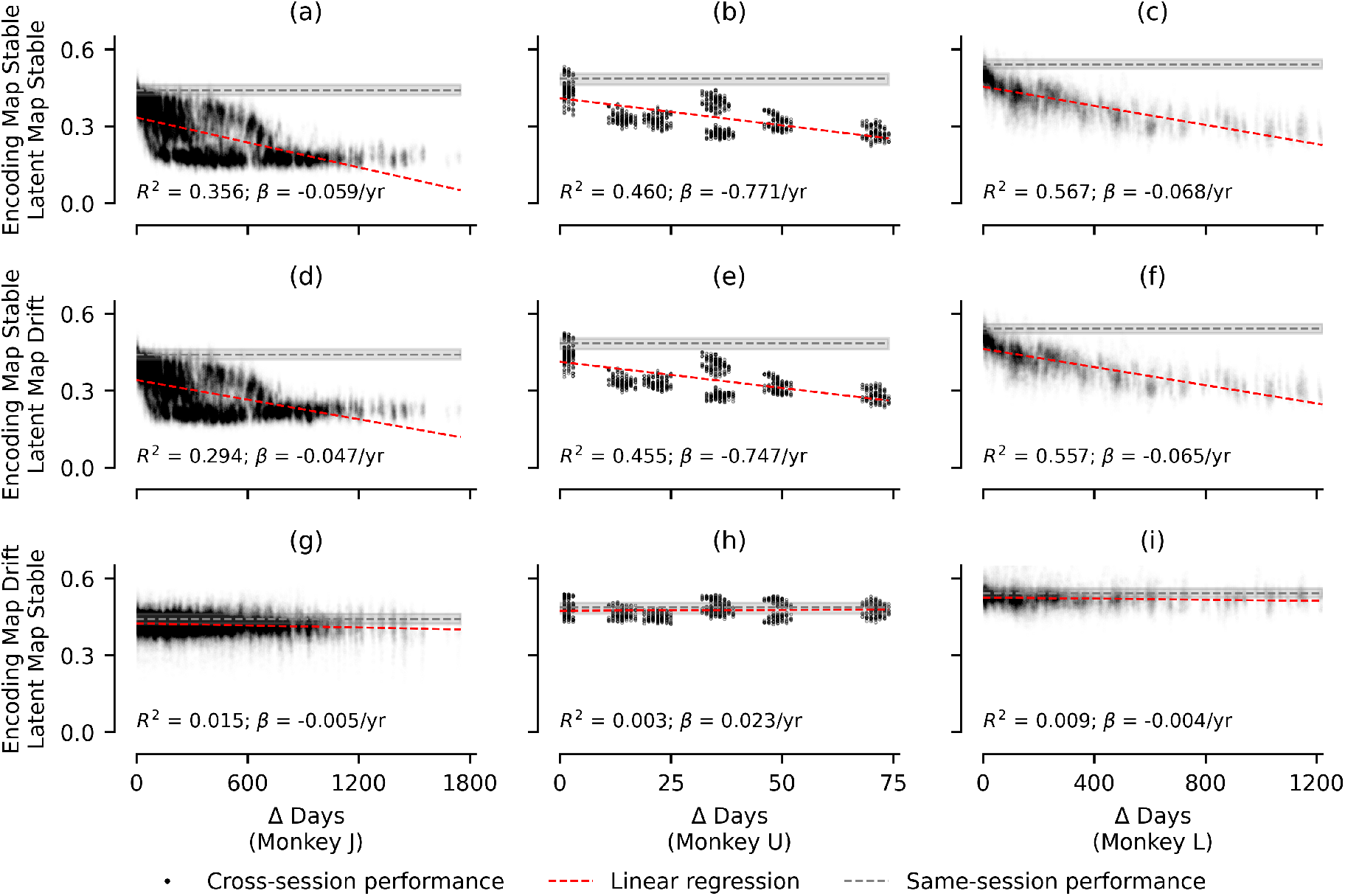
Cross-session decoding accuracy vs. number of days between sessions. Columns indicate results for Monkeys *J* (a, d, g), *U* (b, e, h), and *L* (c, f, i). Black dots indicate decoding accuracy for each pair of experimental sessions *i* ≠ *j* from Supplementary Table 2 using 2-dimensional LDA subspace. Red-dotted line indicates linear regression for performance loss over time. Corresponding *R*^2^ value and slope *β* is included in each subfigure. Grey-dotted line indicates median single-session performance (where both encoding and latent maps are trained and evaluated on data from the same experimental session); shaded area indicates interquartile range. Rows indicate the three hypotheses tested: (a, b, c) stable encoding and latent maps, which resulted in the greatest performance loss over time across all three Monkeys; (d, e, f) stable encoding maps with drifting latent maps, which slightly improved performance over time; and, (g, h, i) drifting encoding maps, alignment rotation, and stable latent map, which achieved stable decoding over months and years of experimental sessions. Results for cursor controller task are shown in Figure 4.

**Figure 4:**
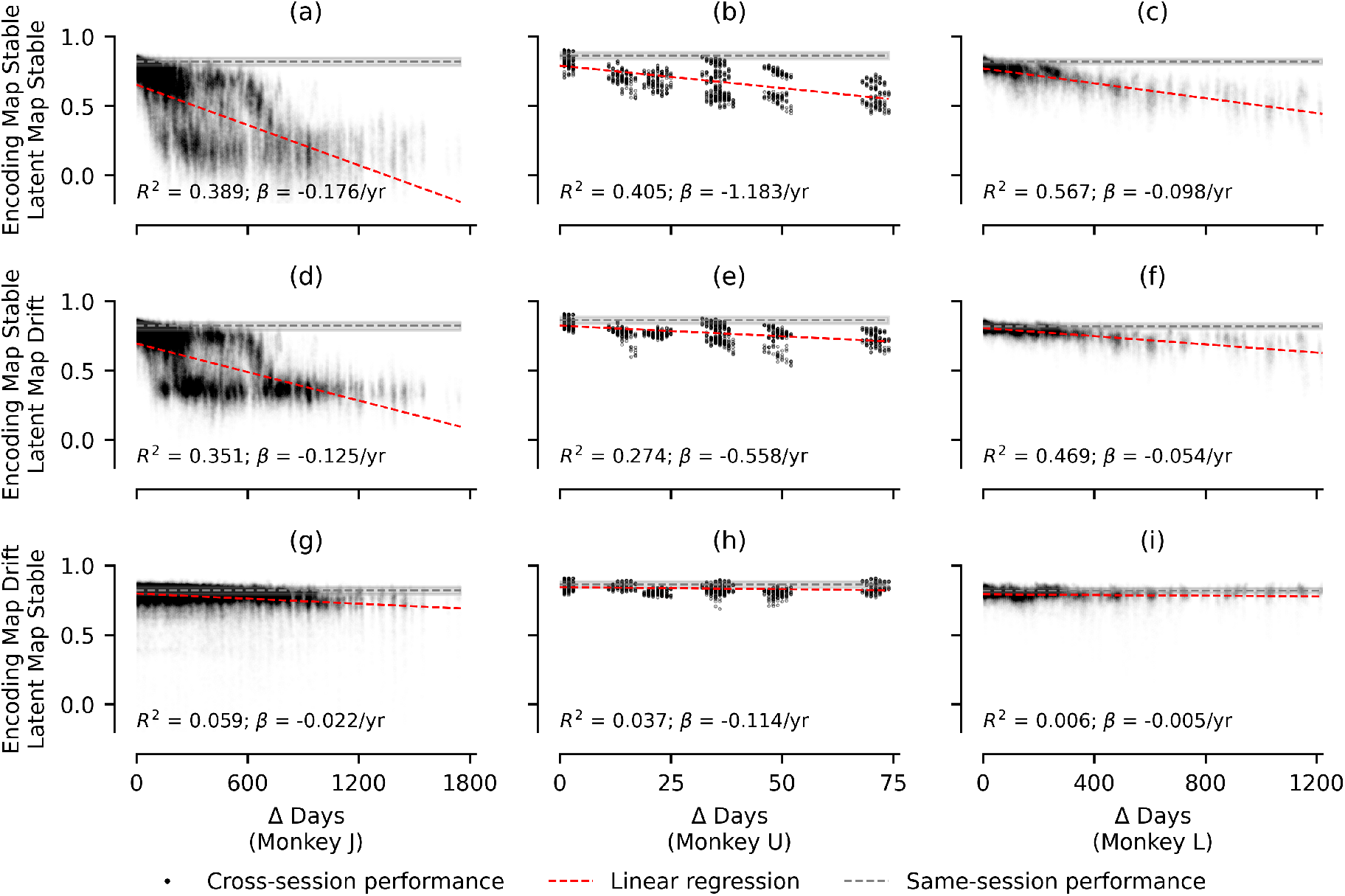
Cross-session cursor position *R*^2^ using controller vs. number of days between sessions. Same as Figure 3, except *y*-axis of each subplot depicts the performance of a linear control system trained on Day *i* and evaluated on Day *j* by comparing coefficient of determination (*R*^2^) between true and reconstructed kinematics.

Both decoding accuracy and controller *R*^2^ values across different experimental sessions decreased drastically as the number of days between experimental sessions increased when using a model with static encoding and latent maps (Figures 3 & 4, Row 1). Fixing only the encoding map and simply re-training the latent map each day only slightly improved model performance over time (Row 2). This indicates that the identified latent space gradually retained less useful information over time. Finally, we found that retraining the encoding map and performing a rotation in low-dimensional space maintained stable performance over time, even with a fixed latent map (Row 3). In fact, this framework’s performance was near-identical to the single-session performance from the previous section (dotted, gray), which is equivalent to retraining both encoding and latent maps every single day. Thus, when the “correct” latent space was identified, the resulting latent trajectories were sufficiently preserved over months and years to allow a static latent map to achieve stable performance for both decoder and controller tasks. Alignment of 2-dimensional latent spaces between sessions was done via rotation mapping; the magnitudes of the required rotation over time are depicted in Supplementary Figure 10, and did not seem to follow any specific trend.

A linear regression was fit for all scatter plots in Figures 3 & 4. This revealed a slope (*β*) describing the rate of change of model performance between sessions, with respect to the number of days between sessions. Loss in model performance over time was an order of magnitude larger for the hypotheses assuming a fixed encoding map. Drift rates of performances across all Monkeys and tasks are compiled in Supplementary Table 3.

### Comparison to previous work

In Gallego, et al. (2020) ^39^, a 10-dimensional encoding map was identified using PCA (unsupervised), and aligned between experimental sessions using CCA (supervised). This achieved stable correlations to hand velocity in a center-out reaching task when using a fixed latent map. This finding was interpreted as evidence that the underlying neural manifold, and in turn the corresponding motor circuits, were roughly 10-dimensional ^39,44^. To study the role of latent space dimensionality, we evaluate long-term performance using three methods for achieving stable decoding and control:

1. The method proposed in this work, which consists of identifying a *k*-dimensional encoding map (or linear subspace) with LDA, applying an optimal alignment rotation *R* ∈ *O*(*k*), and utilizing a fixed latent map. Long-term stability is discussed in the previous section.
2. A modified version of the method proposed in Gallego, et al. (2020) ^39^, which first projects data to a 10-dimensional latent space using PCA, and then performs alignment between sessions by applying CCA to project data to a new, aligned *k*-dimensional latent space.
3. The original method proposed in Gallego, et al. (2020) ^39^, which projects data to a *k*-dimensional latent space using PCA, and then performs alignment between sessions by applying CCA without further dimensionality reduction.

Note that the final two methods are equivalent for *k* = 10 (black-dotted), which are the primary results discussed in Gallego, et al. (2020) ^39^. We apply these methods and track cross-session performance for the decoder and controller tasks, which are depicted in Figure 5. We also include results for decoding reach directions from an entire trajectory and hand-velocity decoding, as in previous literature, in Supplementary Figure 9.

**Figure 5:**
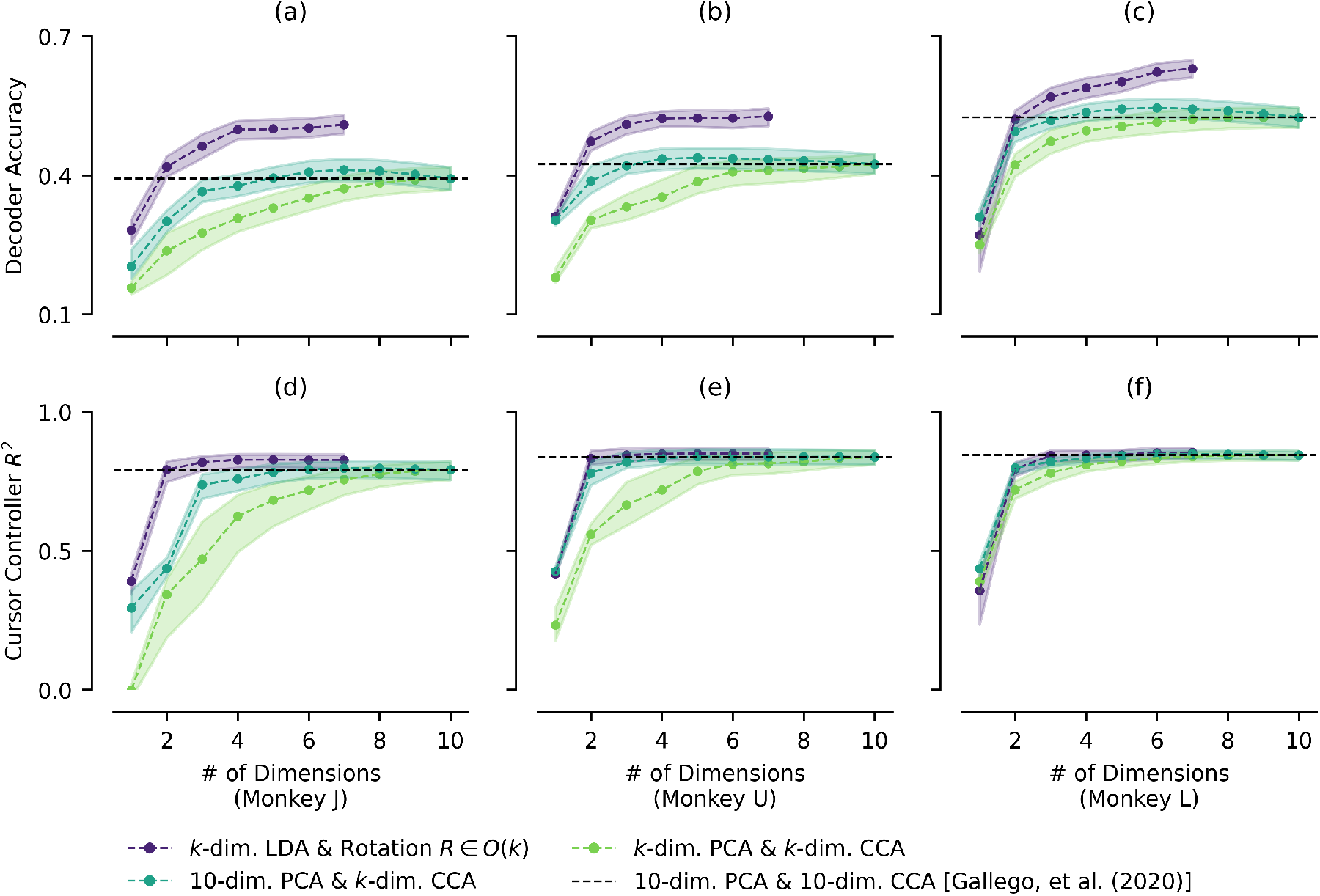
Cross-session performance vs. subspace dimensionality using frameworks for stable long-term performance. As in previous figures, columns indicate results for Monkeys *J* (a, d), *U* (b, e), and *L* (c, f). Rows indicate decoding accuracy (a, b, c) and coefficient of determination between true and reconstructed cursor positions (d, e, f) when trained and evaluated on different experimental sessions. Since all considered methods achieve long-term stability, performances for each method and dimensionality are summarized with a single distribution. Points and dotted-lines indicate median performances for a specific method and dimensionality; shaded regions indicate inter-quartile range. Three frameworks are compared for each subspace dimensionality *k*: *k*-dimensional LDA followed by an alignment rotation *R* ∈ *O*(*k*) (purple), the method proposed in this work; 10-dimensional PCA followed by alignment with *k*-dimensional CCA (teal), a slightly modified version of the framework from Gallego, et al. (2020) ^39^; and *k*-dimensional PCA followed by alignment with *k*-dimensional CCA (light green), the framework proposed in Gallego, et al. (2020) ^39^. The black-dotted line indicates median performance for the last two methods with *k* = 10, the assumed “correct” dimensionality of M1 circuitry. Using LDA with *k* = 2-dimensions achieves and often surpasses this performance in cross-session decoding and control tasks. Data is same as in Figures 3 & 4.

In Gallego, et al. (2020) ^39^, it was argued that the precise dimensionality of the neural manifold was arbitrary, and that estimates of *k* = 6, 8, 10, or 12 all achieved similar long-term performance. Our results corroborate this, as the performance of the model from Gallego, et al. (2020) ^39^ (light green) does plateau around 6–10 dimensions. However, we observe that even when latent space dimensionality for the model proposed in this work (purple) is constrained to the theoretical lower-bound of *k* = 2, its performance becomes comparable to the model from Gallego, et al. (2020) ^39^ with *k* = 10 (dotted-black line). This is most likely because unsupervised methods are not identifying the latent space which regresses against task parameters. Thus, when PCA is used to identify the encoding map, it takes roughly 10 dimensions to capture the latent variables that LDA identifies in only 2 dimensions. This is further supported by the fact that first projecting onto a 10-dimensional space with PCA and then to a *k*-dimensional space with CCA (teal) performs better than the original algorithm in Gallego, et al. (2020) ^39^ (light green) for low values of *k*. The 10-dimensional space identified by PCA envelopes much of the 2-dimensional space spanned by the “correct” latent variables, and a supervised method like CCA can further refine the selection. Thus, the identification of the underlying neural manifold as 6 − 12 dimensional ^39^ might arise from the choice of dimensionality reduction method, rather than the intrinsic neural circuitry (as cross-session stability in performance can be accounted for using only the 2-dimensional latent space identified via LDA.)

### Quantifying drift

In the previous section, we depicted how cross-session performance changes over time; here, we investigate how performance changes with respect to geodesic distance. Since a low-dimensional rotation is used to align latent spaces, this metric must remain invariant when encoding maps are rotated. Thus, rather than quantifying distances between the specific orthonormal basis identified by the encoding map, we quantify distances between corresponding linear subspaces spanned by the orthonormal basis (which is rotation-invariant). This was done using *geodesic distance* between linear subspaces, specifically the implementation derived in Batzies, et al. (2015) ^53^. For further discussion on the interpretation of linear subspaces and geodesic distances in the context of neural data, see the Methods.

In Figure 6, cross-session performance of a model where the encoding map is held fixed and the latent map is re-trained is shown with respect to the squared geodesic distance between the 2-dimensional subspaces extracted by LDA on both days. The gradual decrease in classification accuracy and coefficient of determination are both strongly correlated to geodesic distance. This is further evidence that changes in the relevant subspace (in particular, the subspace identified via LDA) are the dominant contributor to loss of performance of BMI models over time, rather than changes of trajectories in the latent space. Alternative evaluation criteria for decode and control performance are depicted in Supplementary Figure 11. Similar results when PCA was used for the encoding map are shown in Supplementary Figures 12, 13, 14, & 15. Additionally, the gradual increase in geodesic distance between identified LDA subspaces over time is depicted in Supplementary Figure 16.

**Figure 6:**
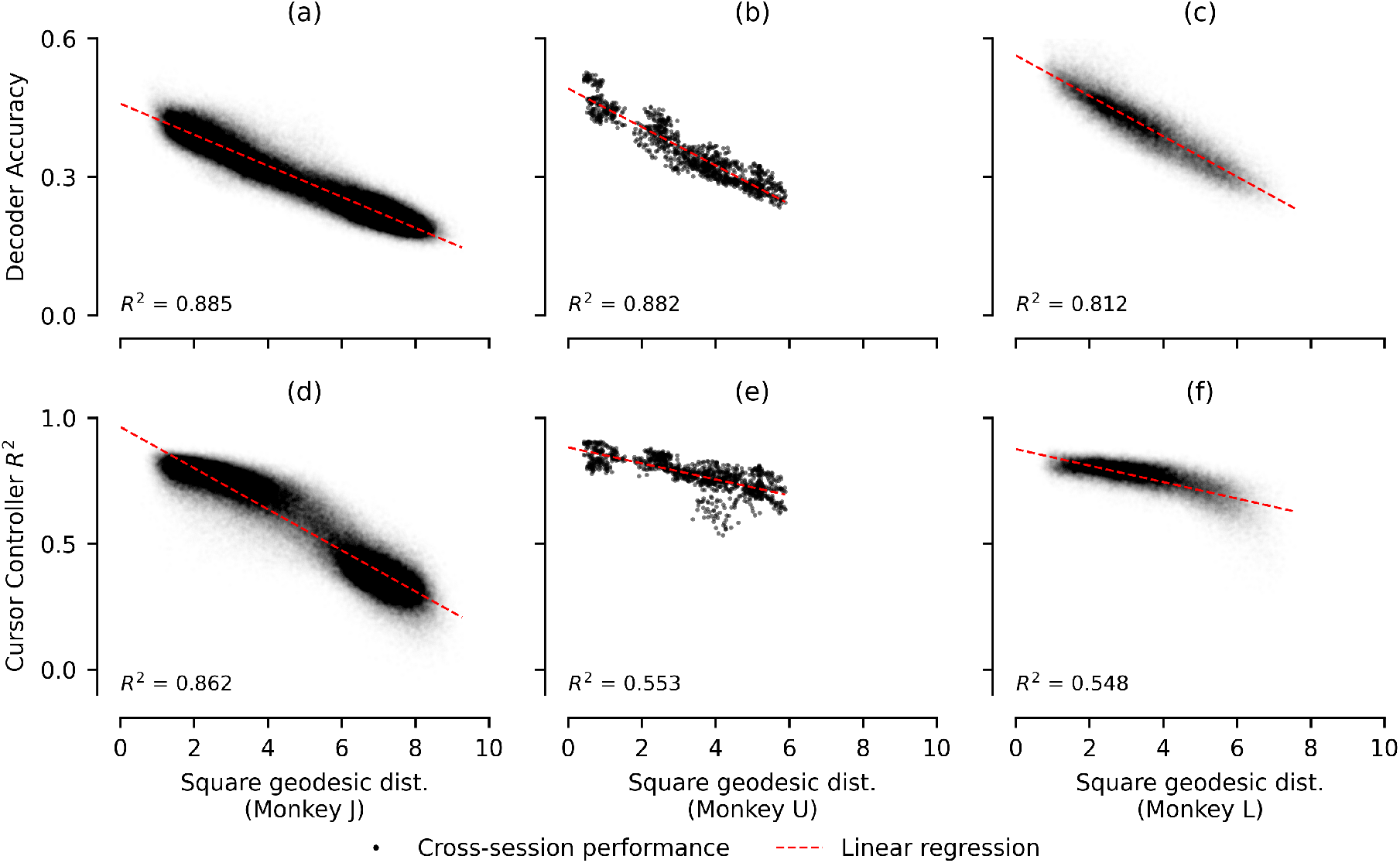
Cross-session performance vs. geodesic distance between LDA subspaces between sessions. Columns indicate results for Monkeys *J* (a, d), *U* (b, e), and *L* (c, f). *Y*-axes for rows indicate decoding accuracy (a, b, c), and coefficient of determination between true and reconstructed cursor positions (d, e, f), when trained and evaluated on different experimental sessions, while keeping the encoding map fixed and re-training the latent map. *X*-axis indicates square geodesic distance between 2-dimensional LDA subspaces identified on Days *i* ≠ *j*, with corresponding *R*^2^ values listed. For both decode and control tasks, loss in performance was strongly correlated to geodesic distances between LDA subspaces. Experimental data same as in Figure 3.

## Discussion

In this work, we proposed a simple framework for extracting an invariant description of motor control, which decomposes the relationship between neural population activity into encoding and latent maps. While extensive research exists on developing latent methods for neural data ^28–33^, this work only considered linear maps, due to their computational efficiency, interpretability, and demonstrated sufficiency for training high-performance BMIs ^37,38^. In general, any encoder-decoder architecture fits into our proposed framework if encoding and latent maps are allowed to be nonlinear. In future work, we hope to address whether our findings hold for nonlinear methods of extracting latent variables from neural data.

We compared how well various linear dimensionality reduction methods performed at extracting behaviorally relevant latent variables by evaluating: (1) accuracy rates for classifying intended reach direction from instantaneous latent states (or alternatively latent trajectories); and, (2) coefficients of determination between true and reconstructed cursor positions using a velocity-based linear control system. While previous literature ^39^ used the coefficient of determination with respect to hand velocity as an evaluation metric, we found that latent states were only weakly correlated to hand velocities (Supplementary Figure 2, Row 4). Regardless of evaluation criteria, the latent variables identified by multiclass LDA consistently outperformed all other linear methods considered in this work. This was despite the LDA subspace accounting for a significantly smaller proportion of the total variance of neural spiking data. While past work has discussed the advantages of using variance explained as a proxy for the utility of the identified subspace ^51^, these results demonstrate that only a fraction of the total variance was required to perform stable decoding of motor behavior in the context of voluntary reaches. Furthermore, performance of the LDA subspace plateaued at roughly 3–4 dimensions for the decoder task, and only 2 for the controller task. Dimensionality of PCA (Supplementary Figures 3 & 4) and dPCA (Supplementary Figures 5 & 6) subspaces had to be substantially increased to achieve similar performance to the 2-dimensional subspace identified by LDA. We suspect this occurs because the subspace relevant for decoder and controller tasks only spans roughly 2-dimensions, and that supervised methods like LDA capture the “correct” latent space in significantly fewer dimensions than other commonly used linear methods. This is somewhat expected, since a simple linear regression against 2-dimensional motor output would span a 2-dimensional subspace. Since the mapping between neural population activity and motor output is most likely nonlinear, the gradual improvement in single-session model performance as dimensionality increases might result from higher-dimensional linear operators approximating nonlinear functions. We must also emphasize a latent model decoding behaviour well *does not imply* that the corresponding latent structure has neurobiological circuit relevance ^54–57^. Significant additional causal interventional studies, such as perturbations ^58^ and knockout studies, are required to help uncover mechanisms ^59^.

Questions regarding the dimensionality of relevant dynamics in motor cortex can be more appropriately investigated when considering datasets spanning multiple sessions over years of experiments. It has been argued that the behaviorally relevant neural manifold in motor regions is roughly 10-dimensional (with various estimates for dimensionality ranging from 5 to 20 ^25,28,36,39,42,43^.) In particular, Gallego, et al. (2020) ^39^ found that 6–12 dimensional subspaces identified by PCA could be aligned between experimental sessions using CCA, allowing for a stable linear map to maintain correlations with hand velocities. The stability of this roughly 10-dimensional latent space was interpreted as a byproduct of the underlying neural circuitry ^39,44^. On the other hand, theoretical work has predicted that the dimensionality of observed dynamics is constrained by task complexity ^47,48,60,61^ and that low-dimensional structure can arise without indicating neural circuit mechanisms ^62^. Here, we investigated three hypotheses concerning long-term stability, under the constraint that the latent space was only 2-dimensional (the theoretical lower bound for a two-dimensional task under a linear model.) Models for decoder and controller tasks were trained and evaluated on datasets from different experimental sessions, and performance was tracked in relation to the number of days between sessions. When all model parameters were fixed, steep performance decay was observed for an offline BMI system as the number of days between sessions increased. When the encoding map (i.e. the latent subspace) was fixed, training a new latent map for each experimental session only slightly improved long-term performance. However, when the encoding map was retrained and identified the “correct” latent space, latent trajectories could be aligned using only a low-dimensional rotation *R* ∈ *O*(2). This allowed a fixed latent map to achieve stable performance in decoder and controller tasks over months and years across all three monkeys considered. These results demonstrate that the 2-dimensional LDA subspace is not only sufficient to infer behavior under task constraints, but also that the corresponding trajectories in this latent space are stable over time. Meanwhile, the set of neural activity patterns regressing to motor behavior (i.e. the encoding map) does change over time. This is most likely due to both recording instability ^39,49^ and non-stationary relationships between neural circuits and behavior (i.e. representational drift ^12,14^). By quantifying this drift using geodesic distance, we demonstrate that changes of the encoding map are a dominant contributor to observed long-term decrease in performance of BMI models ^63–68^. Future work is necessary to differentiate the contributions of electrode instability and encoding drift to the experimentally observed rate of drift of the encoding map, depicted in Supplementary Figure 16. Past work has proposed that the brain balances long-term performance and flexibility by maintaining an invariant model of motor control in the latent space, with a non-stationary neural encoding ^13,16^. Using only longitudinal data from a two-dimensional center-out reaching tasks, it is difficult to distinguish this hypothesis from the alternative, that observed latent stability might arise from constraints imposed by an invariant task topology. However, while the topological argument in the Model Hypothesis section predicts that some endomorphism aligning latent spaces between experimental sessions exists, it does not ensure that only a low-dimensional rotation will be sufficient. In past work, CCA has been used to identify general linear maps that align latent spaces ^39,69^; note that this encompasses both rotation and scaling. To our knowledge, achieving stable long-term decoding without scaling (i.e. aligning latent spaces using only a low-dimensional rotation *R* ∈ *O*(*k*)) has not been reported in previous literature. This potentially indicates that the “power” of the activity patterns regressing to motor behavior in a small recorded neural population remains relatively constant, even when the underlying set of patterns drifts. This also aligns with our observation that the proportion of variance explained by LDA vectors follows a relatively tight distribution, as shown in Figure 2 (Row 2). However, it is also possible that when paired with a simple center-out reaching task, the evaluation metrics (decoder accuracy and coefficient of determination) do not sufficiently vary as latent trajectories are scaled. To better evaluate the direct role that task constraints have on dimensionality and structure of neural manifolds, further work performing longitudinal analysis on datasets of the same monkeys performing tasks with diverse topologies (e.g. a 3D reaching task) is necessary. Additionally, since power and variance explained in a subspace might depend on the number of recording channels, further work is necessary to understand how the number of recorded units affects observed stability.

Accounting for non-stationarity of recorded neural signals in motor regions has been an active area of research, particularly in the domain of brain-machine interfaces ^37,66,70–74^. Stabilizing algorithms that decode behavior from recorded neural activity in an online setting are essential for achieving long-term performance of neuroprosthetic devices. However, evaluating these stabilization algorithms in an offline setting is often unreliable in predicting performance for online applications ^75^. Thus one of the limitations of this work is that it was limited to offline analysis; in future work, we hope to reproduce findings regarding dimensionality and latent stability in online BMI experiments. Nonetheless, offline studies are frequently used to study certain scientific properties, such as how the relationship between neurons, latent dynamics, and behavior either changes or remains stable over time ^39,76^. In Gallego, et al. (2020) ^39^ stable offline decoding performance was achieved by identifying a 10-dimensional neural manifold using PCA (unsupervised), and then aligning manifolds between experimental sessions using CCA (supervised). In this work, we compared how dimensionality of the latent space affected long-term performance of a decoder. We found that when constrained to only 2-dimensions, the LDA-based framework achieved similar performance to the 10-dimensional model discussed in Gallego, et al. (2020) ^39^. When the LDA-based framework was allowed even just a third dimension, it noticeably out-performed the 10-dimensional model. The fact that the LDA subspace explained the observed latent stability using only 2 dimensions has two potential interpretations: (1) that the underlying neural circuitry has significantly fewer degrees of freedom than previously thought; or, (2) that stability of latent trajectories arises from constraints of the 2-dimensional reaching task, not constraints of neural circuitry. We suspect that the latter is more likely, particularly since recent findings indicate that using more flexible models of motor control can reveal up to 200 relevant dimensions in motor regions ^77^. This is not to say that there are no preserved low-dimensional dynamics relevant to motor control, but simply that many experimental paradigms currently used in motor systems neuroscience are not sufficient to distinguish when neural manifolds are exhibiting stability due to preserved task vs. circuit topologies. Previous work has attempted to investigate this alternative possibility; one recent study trained multiple recurrent neural networks (RNNs) to complete a center-out reaching task, while simultaneously constraining them to exhibit different latent dynamics ^46^. This was used to argue that preserved latent dynamics were not a byproduct of networks performing similar motor behaviors. However, this work assumed that the latent space was 10-dimensional and based measurements of dissimilarity in latent dynamics on the entire 10-dimensional latent space. In fact, when CCA was performed to align latent spaces between networks, the two best aligned axes were found to be almost perfectly correlated regardless of how strictly networks were constrained to produce different latent dynamics. Again, statistical techniques revealed a highly-correlated two-dimensional subspace between different networks by virtue of them regressing against identical, two-dimensional behavioral outputs, rather than inherent constraints on network circuitry.

Ultimately, the goal of studying properties of neural manifolds is to gain a deeper understanding of the underlying neural circuitry. In some systems, the topology of low-dimensional neural dynamics has been shown to arise directly from mechanisms at the neural circuit level. For instance, circuit topology and strong local excitatory connections in the head-direction system of fruit flies cause ring attractor dynamics described by neural manifolds ^40,41^. However, the same relationship does not hold in mice, where the circuit mechanisms underlying observed ring attractor dynamics are not well understood ^40^. Thus, while neural manifolds can serve as a link between neural dynamics and circuits, characterizing the neural manifold alone is not sufficient to gain a mechanistic understanding of the underlying circuitry ^40^. In the context of voluntary reaches, we found that using LDA to identify latent variables captured behaviorally relevant low-dimensional dynamics, most likely because the latent space maximized between-class variance (corresponding to the differences in preferred modes of activity during the preparatory phase of reaching ^22,23^) and minimized within-class variance (corresponding to stereotyped trajectories for reaches in the same direction ^24,25^). The LDA subspace not only captured latent trajectories that were preserved over time, it also explained the well-established stability of neural manifolds in motor regions ^39^ in only 2 dimensions. We interpret this as a strong indication that neural manifolds in motor regions are far more constrained by the topology of the motor task; this limits the conclusions that can be drawn about neural circuitry from properties of neural manifolds in motor regions. Thus, we believe that a key emphasis in future studies should be improving experimental paradigms to better evaluate the circuit mechanisms that might give rise to neural manifolds. First, experiments should incorporate motor tasks that enable richer behavioral data acquisition, such as movement in unconstrained environments ^78–82^, complex behaviors, or elements of uncertainty that emphasizes the role of sensory integration. Second, the process of identifying neural manifolds and latent dynamics should be aware of underlying assumptions about neural circuitry ^57^. For instance, the LDA subspace explains motor outputs, but discards behaviorally irrelevant modes of activity and within-class variance between stereotyped trajectories, which might play an essential role if neural circuitry acts as an optimal control system ^83^. Several mechanisms have been proposed for how neural circuitry incorporates sensory information and ultimately drives low-dimensional dynamics in latent space (e.g. stochastic optimal control systems that maximize variance in task-irrelevant dimensions ^83^, balance between inhibitory and excitatory networks in motor circuits ^84^, and active inference ^85^). Testing these mechanisms using the framework of neural manifolds requires mathematical methods for identifying neural manifolds that capture phenomena which might vary depending on circuit architectures, such as preparatory activity, condition-dependent and independent variance, sensorimotor integration, and communication between brain regions, rather than explaining neural variance and task topology. Finally, identification of ground-truth features of neural circuitry should be incorporated into experimental methods. Research in mice has used optical imaging and optogenetic stimulation to record large populations of cortical neurons, in order to identify which statistical properties remain invariant despite representational drift ^12^, and to demonstrate that patterned stimulation of task-relevant neurons can induce the recorded population to generate stereotyped activity patterns correlated to motor behavior ^86^. These advances suggest that optical tools could be used in conjunction with latent variable models to better understand the role of neural circuitry in constraining population dynamics, even in non-human primates, where optogenetic stimulation has been shown to evoke movement in forelimbs ^87^. Frameworks utilizing optogenetic stimulation ^86,88^ and lesioning ^58,89–91^ may also shed light on causal relationships between motor circuits, sensory inputs, and behavioral outputs ^59,88^. Novel experimental paradigms combining optical recordings of population activity, richer descriptions of behavior, sensorimotor integration, and mathematical methods that account for underlying causal mechanisms are necessary to critically evaluate the relationship between preserved latent dynamics, neural manifolds, and motor circuitry.

## Methods

### Data collection

This work conducts a retrospective analysis using data collected from *n*_m_ = 4 rhesus macaques (*J, U, L*, and *O*) performing a center-out reaching task. All animal procedures were reviewed and approved by the Stanford Institutional Animal Care and Use Committee. Hand position for each monkey was tracked at 60Hz using an infrared tracking system (Polaris, Northern Digital Inc, Waterloo, Ontario, Canada). The location of the hand in a straight-forward outreached position was chosen to align with the center of a computer monitor facing the monkey. The hand position relative to this “center” point in space was then projected as a cursor on the computer screen in real-time, allowing the monkey to control the cursor with natural hand motions. Each trial consisted of the following steps:

1. The monkey moves the cursor to an indicated region within a fixed diameter of the center point.
2. A new target appears on the computer screen in one of *C* = 8 positions distributed radially from the center position. (This time indicates the start of the trial).
3. The monkey reaches their hand in the indicated direction to move the cursor into the region indicated by the new target.
4. The monkey maintains the cursor position in the indicated region for a predetermined amount of time (hold time). (See Section on experimental parameters.) If performed successfully, the monkey receives a juice reward.
5. The monkey moves the target back to the center position, and repeats the procedure with another randomly selected radial target for the next trial.

The procedure for aligning kinematics across trials is depicted in Supplementary Figure 17. As shown in Supplementary Figure 17(a), all kinematic data is sampled at a frequency of *F*_*s*_ = 1 kHz with raw cursor positions updated at a frequency of 60 Hz for Monkeys *J, U*, and *L*, and 400 Hz for Monkey *O*, based on hand positions tracked in real time using an infrared cameras. In Supplementary Figure 17(a-e), kinematics are shown in navy blue for *x*- and green for *y*-direction. The trial start (when the new target was randomly selected and shown to the monkey) and end (when the monkey moved the cursor into the indicated target region) are indicated with dashed black lines. A low-pass filter (8^th^-order Butterworth, 20 Hz cutoff, applied forward & backward) was applied, yielding the smoothed data shown in Supplementary Figure 17(b). Cursor velocity and acceleration were approximated by applying a Savitzky-Golay filter (2^nd^-order, 100 ms window) to estimate derivatives of the smoothed cursor position data, depicted in Supplementary Figure 17(c, d) respectively. Normalized velocity and acceleration are obtained by projecting respective vector values onto the target position (via dot product) and scaling to ensure that the maximum value is 1 during the duration of the trial. These are depicted as solid light blue in Supplementary Figure 17(c, d). The point of peak velocity is identified as the sample with maximal normalized velocity that occurs prior to the end of the trial, but more than 100 ms after. The point of peak acceleration is identified as the earliest occurring local maximum of the normalized acceleration occurring between the start of the trial and the point of peak velocity. These points are indicated in light blue in Supplementary Figure 17(c, d), respectively. To extract kinematic samples, the smoothed position data is sampled every 25 ms, from 100 ms before the point of peak acceleration to 400 ms after. These sample times are depicted as the dotted sky-blue lines, with extracted samples shown in Supplementary Figure 17(e).

Neural data is concurrently recorded using a *n*_ch_ = 96-channel microelectrode (Blackrock Neurotech, Salt Lake City, UT) array, implanted in motor regions. Each electrode recorded raw voltage at 30 kHz and identified spikes when the voltage dropped below a constant multiple (in this example, 4.0) of the negative RMS of the recorded signal. An example of extracted spikes for a single channel is depicted as a raster plot in Supplementary Figure 17(f) with black vertical lines indicating a 1 ms sample where a spike was identified. Each sample of kinematic data was paired with a sample of neural data, indicating the number of spikes observed in a given channel in the *t*_bin_ = 25 ms bin preceding the timing of the sample. The extracted sample of neural data is depicted with blue dots, with the corresponding bin width depicted with blue horizontal lines, in Supplementary Figure 17(f).

Thus, each trial consists of *n*_bin_ = 21 total samples of kinematic data (a 2-dimensional vector indicating cursor position) and neural data (an *n*_ch_ = 96-dimensional vector indicating the *state* of the recorded neural population). The neural data consisted of 5 bins preceding and 16 bins following the point of peak acceleration, which was found to usually cover the entire peri-motor period of a reach, spanning a total length of *t*_bin_*n*_bin_ = 525 ms. Given an experimental session with *n*_tr_ successful trials, neural data can thus be described as a matrix 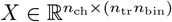. As a final step of preprocessing, to account for any changes in average firing rates within recorded channels, each row of this data matrix was standardized to have mean 0 and variance 1.

### Statistical methods

#### Cross validation

For single-session analysis, a 5 × 2-cross validation ^52^ was performed to estimate the distribution of how well each dimensionality reduction method (or encoding map) performed. Given a dataset from each experimental session, trials were split evenly into training and test sets. The encoding map was identified as the corresponding linear subspace according to each method. The resulting distribution of latent states conditioned on each reach direction was approximated as a Gaussian on the training set and used to train a Naive Bayes decoder (the latent map for the decode task). An affine map transforming latent state to an instantaneous cursor velocity (the latent map for the control task) was also trained to optimally minimize least squares error of cursor position on the training set. The model, consisting of composing the latent and encoding maps to transform neural data into a desired output, was then evaluated on the test set. This process was repeated a total of 5 times with a new randomized train-test split, to estimate the distributions ^52^ of the expected decoding accuracy and controller error. The median and inter-quartile ranges shown in Figure 2 and Supplementary Figures 2, 3, 4, 5, & 6 for each subspace dimensionality are determined from a number of samples equal to 5 times the number of experimental sessions per monkey (*n* = 1190 for *J*, 80 for *U*, 610 for *L*, and 30 for *O*).

For cross-session analysis, each of the 5 test-train splits from each session was paired with a unique test-train split from each of the other sessions. These consisted of the same splits used in the single-session analysis. Thus, the number of data points depicted in Figures 3, 4, & 6, and Supplementary Figures 7, 8, 10, 11, 12, 13, 14, 15, & 16 is equal to 5 times the number of permutations of 2 experimental sessions with no replacement (*n* = 282030 for *J*, 1200 for *U*, and 73810 for *L*).

#### Linear regression

In Figure 6 and Supplementary Figures 11, 12, 14, 13, & 15, regression *R*^2^-values are calculated using a simple linear regression using geodesic distance as the only regressor. In Figures 3 & 4 and Supplementary Figures 7 & 8, regression *R*^2^-values and the rate of change of performance over time were calculated using simple linear regression with the number of days between training and evaluation as the only regressor.

### Encoding maps as neural patterns

Given a neural state 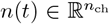, the *encoding map* is defined as a function transforming the neural state to a point in *k*-dimensional latent space. This work focuses on the case where the encoding map is linear, allowing it to be described as a matrix 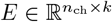. Given the recorded neural trajectory in a single trial 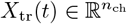, the corresponding trajectory in the latent space is given by

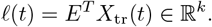

Each element *ℓ*_*i*_(*t*) of the latent state *ℓ*(*t*) describes the temporal evolution of the projection of recorded population activity onto the covariance pattern between channels of the implanted microelectrode array described by the *i*^th^ column 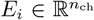 of *E*. Thus, the columns of *E* are referred to as activity patterns, or *encoding patterns*. In the dynamical systems view of motor control, such patterns of neural activity are often modeled to describe neural circuits that ultimately generate motor behaviors ^18^. In this model, the *encoding patterns* considered are the observed patterns of covariance between electrodes that correspond to the projection of neural population activity onto the recording device. The following two restrictions are imposed on the encoding map *E*:

1. For all columns *E*_*i*_ of *E*, ∥*E*_*i*_∥_2_ = 1. This guarantees that for all *i* ≠ *j*, the relative magnitudes of *ℓ*_*i*_ and *ℓ*_*j*_ reflect the relative prominences of patterns *E*_*i*_ and *E*_*j*_.
2. For all *i* ≠ *j*, 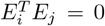. This orthogonality constraint guarantees that the contribution of each pattern to the total variance of the population data is only accounted for once. Thus, reconstructing population dynamics from one latent variable as 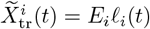 offers no information about another latent variable, since

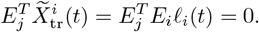

Under these constraints, the set of all possible encoding maps forms the so-called *Stiefel manifold* :

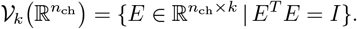

This work is interested in which patterns the brain uses to encode motor control and how these patterns change over time. Consider two consecutive samples of neural population activity, with the relationship:

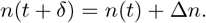

Given an encoding map 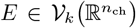 and any orthogonal complement 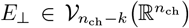, the neural states can be decomposed into the forms

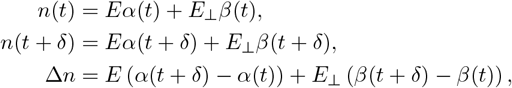

for unique *α*(*t*), *α*(*t* + *δ*) ∈ ℝ^*k*^ and 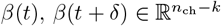. The projections of these states onto the latent space are simply given by *α*(*t*), *α*(*t* + *δ*) since:

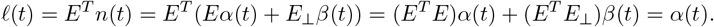

Similarly, *ℓ*(*t* + *δ*) = *α*(*t* + *δ*), implying that the subsequent change in the latent state is given by

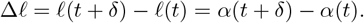

It follows that the two states will have different projections in latent space if and only if *α*(*t*) ≠ *α*(*t* + *δ*). Thus, for the brain to perform motor control by altering the latent state, the change in neural state Δ*n* must have a component that is non-zero when projected onto the latent space under the encoding map *E*. Hence, the equivalence class of neural activity patterns that can be used to perform motor control, are described by the linear subspace given by **span** { *E*_1_, …, *E*_*k*_ }, or for more convenient notation, simply **span** { *E* }. Thus, subspaces spanned by the columns of the encoding map effectively describe the set of neural activity patterns that the brain can use to modulate the latent state and is an object of key interest in this model of motor control.

### Linear subspaces and the Grassmannian

The set of all *k*-dimensional linear subspaces of 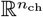 such subspaces is formally defined as the *Grassmannian* manifold. Let ∼ be an equivalence relation on the Stiefel manifold, where

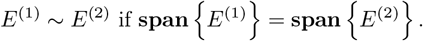

The Grassmannian manifold is often formulated as the quotient manifold generated by this relation:

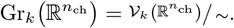

Note that this relation is equivalent to the statement that there exists a matrix *R* ∈ *O*(*k*) such that *E*^(1)^ = *E*^(2)^*R*, where *O*(*k*) is the orthogonal group. Thus, low-dimensional rotations (which are used for aligning latent spaces in this work), do not alter the subspace spanned by the encoding map.

However, this algebraic formulation does not offer a standard matrix representation for every linear subspace. Alternatively, the so-called *isospectral picture of the Grassmannian* ^92^, is an explicit embedding in 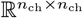:

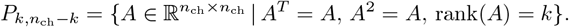

This is diffeomorphic ^92^ (and often treated as equivalent ^93^) to the more common quotient-space formulation of the Grassmannian. Thus, the Stiefel manifold can be formulated as a vector bundle on the Grassmannian, where the map defined *E* ↦ *EE*^*T*^ serves as a convenient numerical implementation of the corresponding surjection 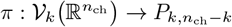 ^93^.

Changes in the set of patterns that the brain uses to perform motor control over time can thus be modeled as a process evolving on the Grassmannian manifold. Since this is a Riemannian manifold, a notion of “distance” can be defined between two samples as the length of the geodesic curve connecting them. Given samples 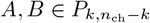, the geodesic distance can be expressed in closed-form:

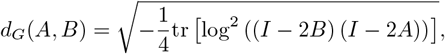

with log indicating the matrix logarithm ^53^. Note that this formula is restricted to 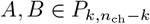 for which the orthogonal matrix (*I* − 2*B*)(*I* − 2*A*) has no negative real eigenvalues, a constraint which was satisfied by all pairs of encoding maps in our dataset.

### Linear dimensionality reduction

Linear methods offer increased model simplicity, interpretability, and convenient mathematical properties. In past work, linear methods have been sufficient for the design of high-performance BMIs ^37^; furthermore, linear dimensionality reduction methods have also been successful at identifying relevant latent variables that improve BMI performance ^38^. Four linear dimensionality reduction methods are considered that are common in the literature. In particular, these are methods that also offer clear reformulations as *subspace methods*, allowing them to be interpreted as sets of activity patterns that fit neatly into the framework discussed in the previous section.

#### Principal component analysis (PCA)

The neural data from an experimental session with *n*_tr_ trials can be described as a matrix 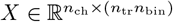. As part of preprocessing, the rows of *X* are z-scored such that each row has mean 0 and variance 1. Thus, the covariance matrix of the recorded channels can be formulated as

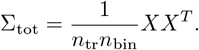

Principal component analysis (PCA) identifies 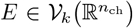 as the matrix whose columns are the eigen-vectors with the *k* largest corresponding eigenvalues. PCA assumes a specific *orientation* of the columns of *E* based on their corresponding eigenvalues; since this is equivalent to the amount of variance explained in the data, referred to as a *varimax rotation*.

However, there is no reason to expect the varimax rotation to orient latent trajectories consistently. Thus, we reformulate PCA as an optimization problem with a *rotation invariant* objective function and address the issue of finding the correct orientation of the columns later:

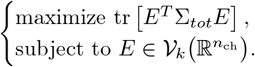

Here, the trace of the matrix depends not on the specific columns of *E* but on the subspace spanned by the columns of *E*. This conveniently fits into our interpretation of linear methods as identifying the space of relevant patterns that the brain utilizes during motor control.

Finally, note that if no spikes were recorded in a channel, the corresponding row of *X* would be 0, resulting in an eigenvalue of Σ_tot_ also being 0. In the next section, it is shown that invertibility is an convenient property for the covariance matrix. Thus, the covariance matrix is typically formulated instead as

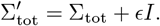

Eigendecomposition of this matrix can be thought of as solving the problem of least squares with regularization ^50^. In this work *ϵ* = 10^−5^ is used to ensure numerical stability in rare cases where no spikes are recorded in certain channels.

#### Canonical correlation analysis (CCA)

While PCA takes a single dataset and finds the subspace with the maximum amount of explained variance, Canonical correlation analysis (CCA) maximizes correlation between datasets. Assume two datasets with the same number of samples are provided, but not necessarily the same number of dimensions:

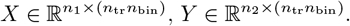

Let Σ_*XX*_, Σ_*Y Y*_ be the total covariance matrices for the datasets *X, Y* respectively, as defined in the previous section. (Regularization can be additionally imposed to ensure these matrices are invertible, as in the previous section). Additionally, 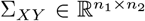 be the associated cross-covariance matrix. CCA seeks to find linear subspaces 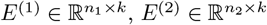 such that the projected datasets (*E*^(1)^)^*T*^ *X* and (*E*^(2)^)^*T*^ *Y* have maximum correlation. This can be calculated by using the singular vector decomposition (SVD) of the whitened cross-covariance matrix 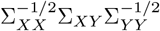. That is, if 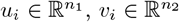 is the left-right singular vector pair with the *i*^th^ largest singular value, then the *i*^th^ columns of *E*^(1)^, *E*^(2)^ are given respectively by 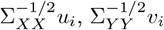^94^.

In this work, CCA is used in two contexts: (1) as in past work ^39^, it is used to identify low-dimensional projections that maximally correlate neural data recorded in different experimental sessions. In this case, both *X* and *Y* are sets of neural data; (2) it is used to identify a linear subspace of neural data from a single session that best regresses against kinematic data. In this case, *X* is a set of neural data, and 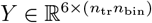 is a corresponding dataset where each sample contains the position, velocity, and acceleration of the cursor on the computer screen in real time. In this case, only the linear subspace of the neural data is relevant; hence, CCA can be reformulated as a subspace method using the following optimization problem:

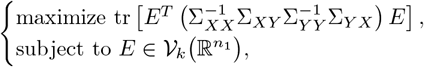

which can be solved via eigendecomposition^95^.

#### Demixed principal component analysis (dPCA)

PCA seeks to explain a maximal amount of total variance, which both condition-dependent and condition-independent sources contribute to ^96^. Since an ideal latent description would allow for different outputs for different task conditions, subspace methods that primarily capture condition-dependent variance are particularly relevant ^51^.

For each reach direction *c* ∈ { 1, …, *C* }, define *µ*_*c*_ as the mean neural state across all bins for all trials reaching in direction *c*. Let 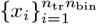 be the columns of the data matrix *X*. The matrix 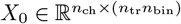 can be defined such that the *i*^th^ column is the mean state *µ*_*c*_ for the reach direction *c* corresponding to the sample *x*_*i*_. Let *A*_OLS_ be the ordinary least squares solution to the problem

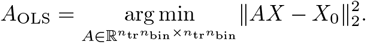

Demixed PCA identifies a map *E* by performing PCA on the projected dataset *A*_OLS_*X*. Note that *A*_OLS_ has a closed-form solution:

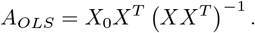

(For numerical stability in cases where *XX*^*T*^ is not invertible, the approximation 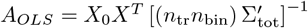 is often used, as discussed in the section on PCA). Thus, the columns of *A*_OLS_ are linear combinations of the columns of *X*_0_, which consist only of mean vectors 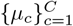. It follows that eigenvectors of that arise from performing PCA on *A*_OLS_*X* with non-zero eigenvalues must span the same linear subspace as the set 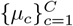. Since each row of the original data matrix *X* had mean 0, there must exist some linear combination of the vectors 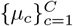 that sums to 0, i.e. the mean states are not linearly independent. Hence, the rank of *A*_*OLS*_*X* (and the maximum number of features that can be extracted through dPCA) is bounded above by *C* − 1. Every dataset considered in this work achieved this upper bound, most likely since the number of channels *d* is much greater than the number of reach directions *C*.

#### A note on condition-dependent variance

Suppose that rather than projecting the true neural data *X* onto the matrix of means *X*_0_ using least squares, the neural state simply took on the constant value *µ*_*c*_ during a reach in direction *c* ∈ {1, …, *C*}. This is equivalent to simply taking *X*_0_ as our data matrix, and serves as an approximation for neural data with all condition-independent sources of variance removed. Assuming there are *n*_*c*_ trials for each reach direction *c* (and therefore 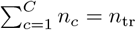), the covariance matrix becomes

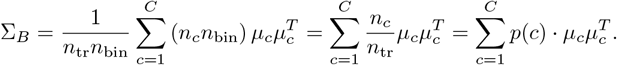

This is equivalent to the formulation of the between-class scatter matrix as described in existing literature ^97^. Thus, performing PCA on this dataset involves performing eigendecomposition on the between-class scatter matrix Σ_*B*_, which yields a set of *C* − 1 eigenvectors with non-zero eigenvalues that span the same subspace as the demixed prinicpal components. This method is closely related to dPCA; across all datasets considered in this work, there was a negligible difference between the projection map identified via dPCA and this method. Hence, dPCA can be approximated by solving the optimization problem:

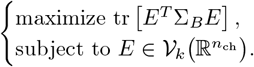

#### Multiclass linear discriminant analysis (LDA)

For each *c* ∈ {1, …, *C*}, let 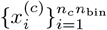 be the set of columns of the data matrix *X* which correspond to a sample of neural activity recorded during a reach in direction *c*. A between-class scatter matrix can therefore be constructed as:

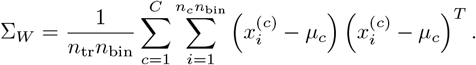

Note that this is essentially a weighted average of the covariance matrices corresponding to individual reach directions. This formulation has the convenient quality that the total covariance is simply the sum of between- and within-class scatter matrices ^50,97^:

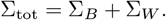

Linear discriminant analysis attempts to find a low-dimensional projection of the data maximizes discrimination between classes, by maximizing between-class variance while minimizing within-class variance. This is done by maximizing the Fisher quotient:

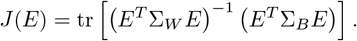

Note that in general, Σ_*W*_ is not guaranteed to be invertible, e.g. the case where a single channel records 0 spikes. To ensure numerical stability in this case, a similar form of regularization as described in the section on dPCA is used by interchanging:

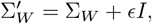

with *ϵ* = 10^−5^ for the analysis in this paper. The objective function *J*(*E*) is maximized by identifying the *k* eigenvectors of the matrix 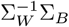 with the largest corresponding eigenvalues. These eigenvectors will be strictly real (with real eigenvalues), with the additional benefit or remaining invariant under changes of the coordinate system (see the discussion in Chapter 10 ^97^). As was the case with dPCA, the number of positive real eigenvalues is bounded above by *C* − 1. Furthermore, while Σ_*W*_ and Σ_*B*_ are both symmetric matrices, 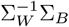 is generally not, meaning that the columns of *E* will not be orthogonal. To address this, Gram-Schmidt orthogonalization is used to project 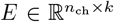 to the Stiefel manifold 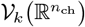. As discussed in section on Grassmannian manifolds, the subspace spanned by the columns of *E* (which does not change under Gram-Schmidt) is an object of key interest, rather than the choice of the specific columns forming *E*. Hence, this heuristic method is satisfactory for our purposes. In the following sections, other methods for performing discriminant analysis with an explicit orthogonality constraint are discussed. Ultimately, when applied to the data in this work, it was found that LDA as described had the best performance at both decode and control tasks. Thus, the performance of the methods in the following sections is not covered in this work. However, the discussion of similar methods helps build an intuition for how such methods relate to one another.

#### Trace-ratio LDA (trLDA)

While the objective function for ordinary LDA seeks to maximize the trace of a matrix ratio, alternative formulations of linear discriminant analysis have sought to instead maximize the ratio of traces:

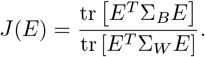

When paired with an additional orthogonality constraint 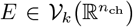, this problem can be referred to as trace-ratio LDA (trLDA), but note that a substantial body of literature exists on this problem under the name Foley-Sammon LDA (after the seminal work by Foley and Sammon ^98^). Unlike the formulation of ordinary LDA, this formulation is not invariant to a specific coordinate system ^97^, which can be a useful property in the analysis of neural data. Past work has suggested solving this optimization problem by either iteratively identifying columns of *E* ^99^, or a general optimization software than can act directly on the Stiefel manifold ^100^ such as *manopt* ^101–103^. In this work, an iterative algorithm shown in Algorithm 1 was utilized instead. Global convergence properties of this algorithm have been discussed in past work ^104^. The key insight of this scheme is the relationship of the trace-ratio problem to the trace-difference problem discussed in the next section.

##### Algorithm 1

Trace-Ratio LDA Global Maximum

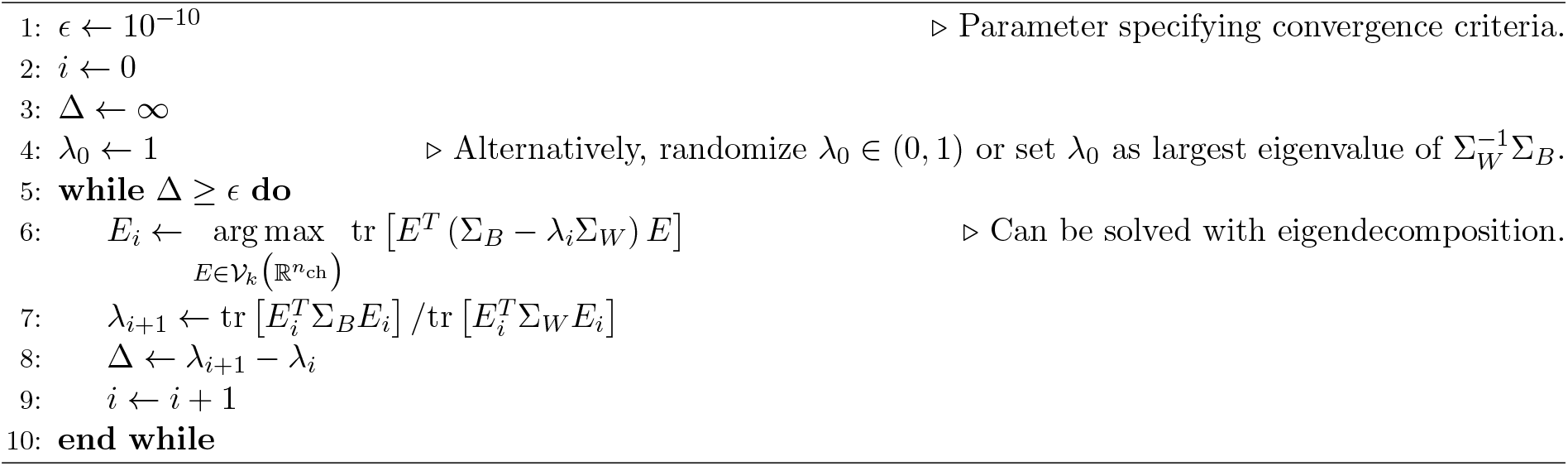

#### Trace-difference LDA (tdLDA)

The final form of LDA considered is what we refer to trace-difference LDA, or tdLDA. As the name implies, this method involves solving the following optimization problem:

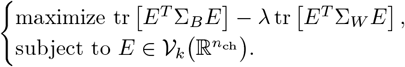

This method involves a tunable hyperparameter *λ* > 0, and can be solved using eigendecomposition of the matrix (Σ_*B*_ − *λ*Σ_*W*_), since this matrix is symmetric and will have orthogonal eigenvectors. It follows from the Algorithm 1 that this formulation is equivalent to trace-ratio LDA in the case where

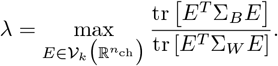

Furthermore, note from the analysis in the section on dPCA that this method will converge to an approximation of demixed PCA as *λ* → 0. On the other hand, as *λ*→ ∞, the solution of the optimization problem simply becomes the projection that minimizes within-class variance. In the case of neural data, this was often simply the null-space of the data (if such vectors existed), or simply the principal components with the smallest corresponding eigenvalues. Hence, the parameter *λ* can be thought of as a regularization constant which decreases bias towards large principal components as it increases.

#### A note on LDA and dPCA

Out of all the methods of deriving a discriminating subspace, simply performing LDA using the Fisher Quotient followed by Gram-Schmidt orthogonalization yielded the highest performance for both decoding and control tasks in the single-session case. While this method for LDA is distinct from the other proposed methods, it is deeply related. It can be approximated as trace-difference LDA with an appropriate choice of *λ >* 0. As discussed in the previous section, choosing *λ* = 0 also serves an approximation of dPCA. Thus, LDA can be approximated as a version of dPCA with additional regularization that removes bias towards large sources of within-class, or alternatively condition-independent, variance. In practice, it was found that these tended to be the largest principal components. By removing this bias, LDA performed better at isolating sources of condition-dependent variance than dPCA, i.e. identifying the subspace relevant to differentiating between task conditions in the lowest number of dimensions possible.

### Gaussian Naive Bayes decoders

#### Classifying a sample

Let 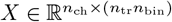 be a dataset where each column indicates a sample bin of neural population activity from one of *n*_tr_ center-out reaching trials. As in the section on LDA, for each reach direction *c* ∈ {1, …, *C*}, let *n*_*c*_ indicate the number of trials and let 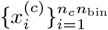 be the set of columns of the data matrix *X* corresponding to reaches in direction *c*. Under an encoding map 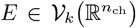, the conditional mean and covariance of the projected dataset corresponding to each reach direction is given by

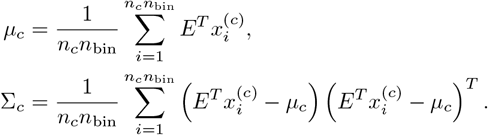

Let *p*_*c*_ : ℝ^*k*^ → ℝ_+_ be the probability density function corresponding to a multivariate normal distribution 𝒩 (*µ*_*c*_, Σ_*c*_):

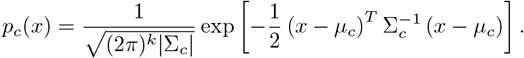

Under a Gaussian Naive Bayes model, the latent map can be thought of as a classifier *L* : ℝ^*k*^→ { 1, …, *C* }which uses Bayes rule to choose a reach direction with the highest conditional probability given an observation ^50^. Given a sample of neural population activity 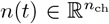, this model would find the projection to the latent space *ℓ* = *E*^*T*^ *n*(*t*) given by the encoding map, and then apply the latent map:

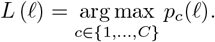

#### Classifying a trajectory

The simplest possible model assumes that within a neural trajectory, each sample is an independent observation. This is trivially false in the case where any latent dynamics affect the trajectory, but yields a convenient mathematical solution for performing classification efficiently. Given a trajectory in the latent space consisting of samples 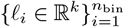, the latent map *L* : ℝ^*k*^ *×* {1, …, *n*_bin_} → {1, …, *C*} is given by:

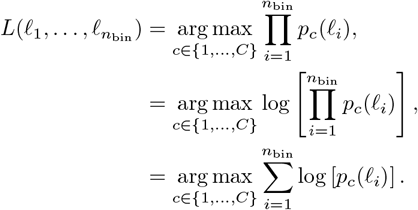

### Optimal linear control

Consider a single trial in the given motor task, with samples of kinematic and neural data collected according to the procedure in the section on data collection at times 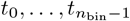. Let 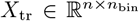 be the *n*-dimensional samples of neural population data collected for this trial, where each entry denotes the number of spikes observed in the preceding *t*_bin_ = 25 ms bin. Under an encoding map 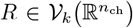, the latent trajectory is given by

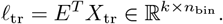

An affine latent map can be modeled as a matrix *L* ∈ ℝ^(*k*+1)*×*2^, transforming the latent trajectory into an estimated velocity trace given by:

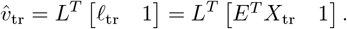

If the true neural trajectory takes the form of a piecewise linear function between the sampled values, the resulting estimated estimated velocity trace is also given by a piecewise linear function. This allows the resulting displacement (corresponding to the sample times) to be expressed using the integrator matrix:

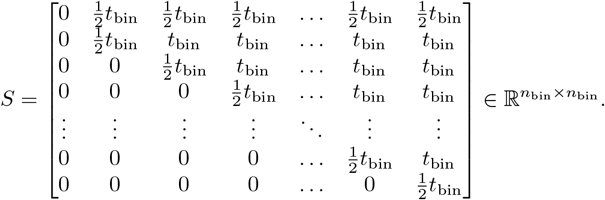

Let the true cursor position be given by the vector

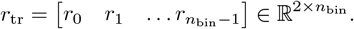

Assuming that both true and reconstructed cursor positions begin in the same position, the reconstructed cursor position can be expressed

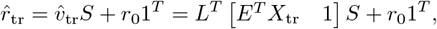

where 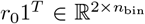 is the matrix where each column is the initial position *r*_0_ ∈ ℝ^2^. Since the residual difference between true and approximated cursor position depends linearly on *L*, the latent map can be trained using least squares. To quantify how close the true and reconstructed cursor trajectories are, we use three evaluation metrics. First, we use the coefficient of determination (*R*^2^). Let 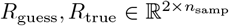 be the reconstructed and true cursor positions for the entire dataset, and *R*_resid_ = *R*_true_ − *R*_guess_ the matrix of residuals. Let Σ_resid_, Σ_true_ ∈ ℝ^2*×*2^ be the covariance matrices of *R*_resid_ and *R*_true_, respectively. Then we calculate the coefficient of determination using

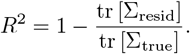

(This is the same formula used to determine *R*^2^ between neural state and hand velocity, which is trained using an affine map.) Alternatively, we also consider the average Euclidean distance between each reconstructed and true cursor position. If the reconstructed cursor position for a given trial is described by the vector

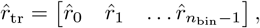

then the average *l*^2^-norm error for a trial is given by:

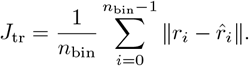

Similarly, the average *l*^2^-norm error is defined for an entire experimental session by averaging the error obtained for each trial. Finally, note that training the latent map for the control task involves identification of the matrix *L*.

### Optimal alignment rotation

Let 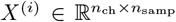 be the neural data collected from an experimental session on Day *i*. Let 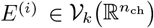 be the corresponding encoding map identified through one of the subspace methods discussed previously. Let *n*_*c*_ indicate the number of total trials with the target placed in radial direction *c* ∈ {1, …, *C*}.

The total number of trials is thus

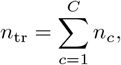

with *n*_samp_ = *n*_tr_*n*_bin_. For each reach direction *c* ∈ { 1, …, *C* }, let the neural trajectories corresponding to each individual trial be given by:

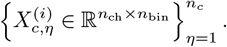

The average latent trajectory for each reach direction is thus given by:

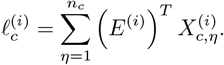

Thus, regardless of the number of trials achieved in a given experimental session, the matrix of average latent trajectories can be expressed as:

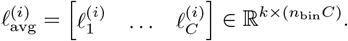

Given two such matrices 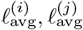 obtained from different experimental sessions, obtaining the optimal alignment rotation becomes an optimization problem:

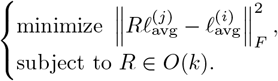

This has a closed-form solution. Let

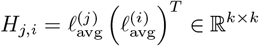

be the cross-covariance matrix. Let the singular value decomposition (SVD) of *H*_*j,i*_ be given by:

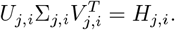

Then the rotation that solves the optimization problem above is given by:

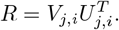

Note that this method for finding an optimal rotation does not depend on the number of trials completed in a session, or the order in which the trials occur, which gives it a significant advantage over methods for aligning latent spaces proposed in past literature ^39^.

### A note on alignment rotations

While each linear method outputs a specific orthonormal basis, the choice of basis depends on some underlying eigenspectrum, and there is no reason to expect the basis to consistently have the same correlation to movement parameters. Thus, each method is instead treated as a *subspace method*. The linear subspace spanned by the columns of *E* is of key interest, as these are the set of activity patterns the brain can utilize to modulate the latent state and perform motor control under this model, rather than simply the exact patterns making up each column of the encoding map *E*. Note that applying an arbitrary rotation in low-dimensional latent space does not change this subspace, since

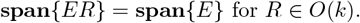

Thus, the choice of using only a low-dimensional rotation to align trajectories in latent space is consequential. For instance, using an affine transformation would no longer yield a clear interpretation of encoding maps as activity patterns; using any linear map would cause the shape of Gaussian distributions in the latent space to change, which limits conclusions that can be made based on the performance of the Gaussian Naive Bayes decoder. The fact that only a rotation is necessary to consistently pin the latent space to behaviorally relevant axes indicates that the identified subspace has contains an invariant latent trajectories that encode motor control in a consistent way.

Furthermore, some degree of latent alignment is always necessary, since the encoding maps are found using eigendecomposition based methods, which do not have a positive or negative orientation by default. For instance, given identical data on Days *i* and *j*, the resulting encoding maps could very well differ by a multiple of − 1 for each column of *E*. This would cause a latent map trained on Day *i* to perform poorly on Day *j*, despite the underlying data being identical. Thus, at the very least all 2^*k*^ encoding maps resulting from multiplying columns of *E* by − 1 must be considered to say whether a latent description is invariant. This set forms a subset of *O*(*k*). Using a rotation generalizes this edge case by considering all possible bases spanning the same subspace, allowing the extent to which latent trajectories are preserved within a linear subspace (or intrinsic manifold ^42^) to be studied.

### Experimental parameters

Data for Monkeys *J* and *L* was collected over years of experiments across multiple studies from the Neural Prosthetic Systems Lab (NPSL). Data collection was in conjunction with brain-machine interface (BMI) experiments. All datasets consisted of a single, contiguous block of reaching trials during a single day’s experimental session, with BMI control reaching trials typically performed in a separate contiguous block of trials afterwards. No neural data from trials utilizing BMI control, or interleaved with trials using BMI control, was included in this study.

Data from Monkey *U* was collected as part of a set of studies from the Brain Interfacing Lab that performed electrolytic lesions that preserved the microelectrode array ^89,105^. Prior to each of the four lesions, four days of “pre-lesion” experiments were performed, which make up the 16 total experimental sessions for this monkey.

Data from Monkey *O* was collected during training to perform more complicated motor behaviors for ongoing studies from the Brain Interfacing Lab. Experimental parameters for all monkeys are summarized in Supplementary Table 4.

## Data Availability

Data will be made publicly available at the Stanford Digital Repository, with deposition currently pending.

## Code Availability

All computational methods are described in full detail in the Methods section. Code will be made publicly available at the Stanford Digital Repository, with deposition currently pending.

## Acknowledgments

The members of the Brain Interfacing Laboratory are Michele Wechsler, Alexandra Paraskevopoulou, Mackenzie Risch, Stephen I Ryu, Iliana Bray, Alissa Ling, Michael Silvernagel, Kenji Marshall, Alice Tor, Yuxin Wu, and Sydney Hunt. We thank Kimberly Chin for administrative support. We thank the members of the Neural Prosthetics Systems Laboratory, in particular Jonathan Kao and Sergey Stavisky for data collection of certain datasets for Monkeys J and L.

## Author Contributions

MUA designed study framework, reviewed and extracted relevant datasets, programmed data processing pipelines, developed mathematical framework for modeling data, analyzed data, prepared figures, and wrote original paper (conceptualization, data curation, formal analysis, methodology, software, visualization, writing – original draft). SEC led data collection in Monkey *U* and supported data collection in Monkey *O* (data curation, investigation). EJJ led data collection in Monkey *O* (data curation, investigation). PN contributed to all aspects of the research (conceptualization, data curation, funding acquisition, investigation, methodology, project administration, resources, supervision, validation). All authors reviewed and edited the paper (writing – review & editing). All members of the Brain Interfacing Lab also provided insight on review and editing.

## Ethics Statement

All procedures were reviewed and approved by the Stanford Institutional Animal Care and Use Committee.

## Supplementary Information

**Supplementary Table 1:** Summary of past estimates and models of dimensionality of motor regions.

| Paper | Dimensionality | Additional context |
| --- | --- | --- |
| Churchland, et al. (2007)<sup>7</sup> | identified 10 or 29 | depended on choice of method |
| Yu, et al. (2009)<sup>28</sup> | identified 10 |  |
| Churchland, et al. (2012)<sup>25</sup> | modeled as 6 | oscillatory dynamics spanned 2 |
| Sadtler, et al. (2014)<sup>42</sup> | modeled as 10 |  |
| Gallego, et al. (2018)<sup>36</sup> | modeled as 8–15 | called exact number arbitrary |
| Gallego, et al. (2020)<sup>39</sup> | modeled as 6–12 |  |
| Sabatini, et al. (2024)<sup>43</sup> | identified 5–9 or 6–20 | depended on choice of method |

**Supplementary Table 2:**
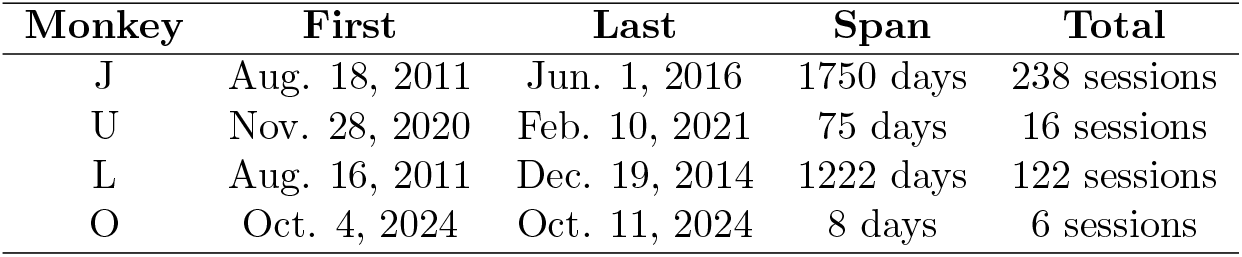
Summary of experimental sessions per monkey. Columns indicate dates of first and last experiments, span of period of experimentation, and total number of sessions. All four monkeys were used for the single-session analysis. Only three monkeys with multiple weeks of data (*J, U*, & *L*) were used for the long-term cross-session analysis, as data for Monkey *O* only spanned eight total days. In Figure 2, each point represents a distribution consisting of a number of samples equivalent to five times the number of experimental sessions, due to the choice of cross-validation ^52^: *J*, 1190 samples; *U*, 80 samples; *L*, 610 samples; *O*, 610 samples. Cross-session results depicted in Figures 3, 4, 5, & 6 compare each of these partitions to a corresponding partition from a different session, resulting in a total of: *J*, 282030 samples; *U*, 1200 samples; *L*, 73810 samples.

| Monkey | First | Last | Span | Total |
| --- | --- | --- | --- | --- |
| J | Aug. 18, 2011 | Jun. 1, 2016 | 1750 days | 238 sessions |
| U | Nov. 28, 2020 | Feb. 10, 2021 | 75 days | 16 sessions |
| L | Aug. 16, 2011 | Dec. 19, 2014 | 1222 days | 122 sessions |
| O | Oct. 4, 2024 | Oct. 11, 2024 | 8 days | 6 sessions |

**Supplementary Fig. 1:**
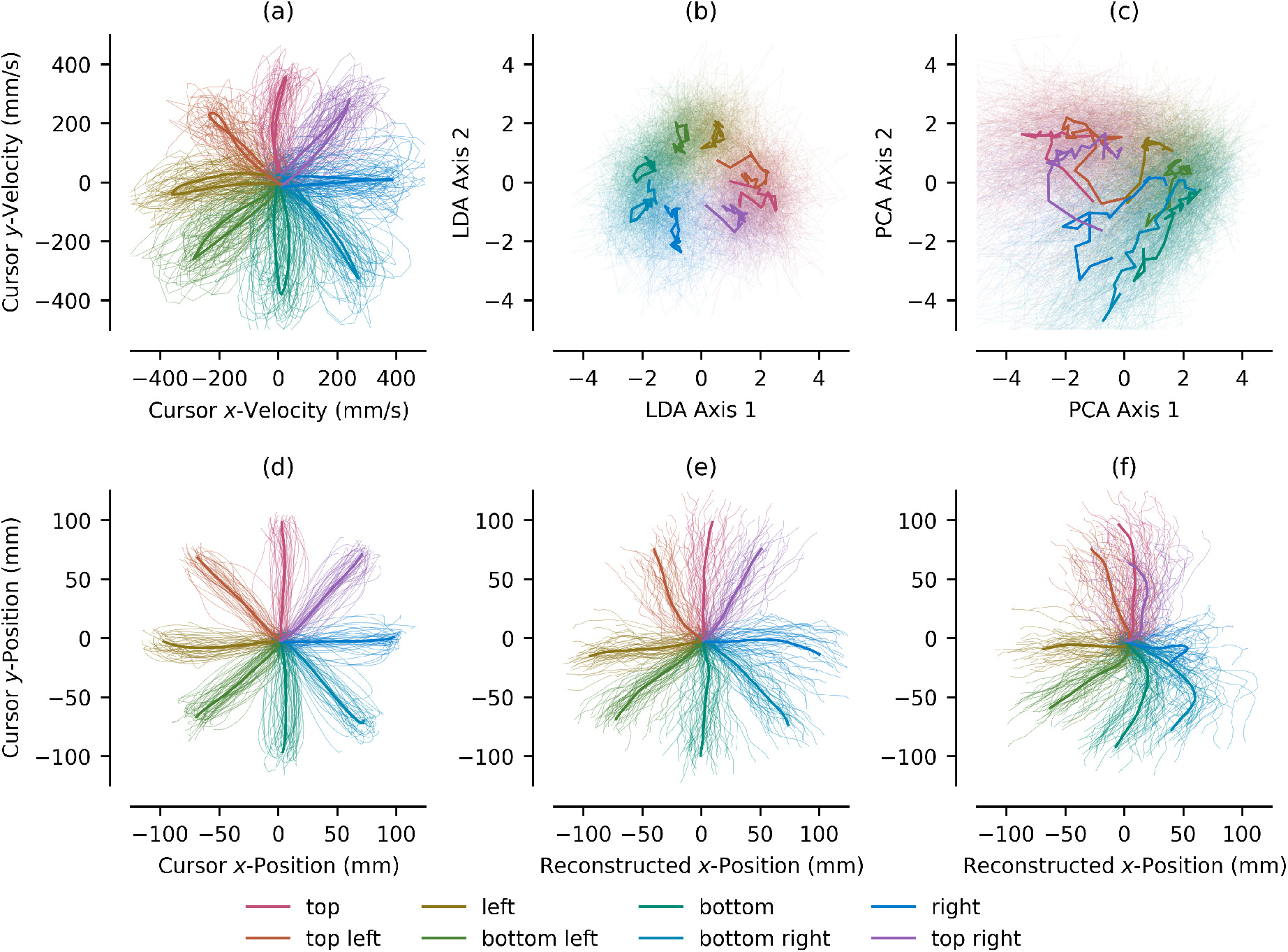
Kinematic and neural trajectories from experimental session U201201 01. Colors indicate trajectories for each of eight target directions in the radial center-out reaching task. Thicker lines indicate average trajectories for each task condition, thinner transparent lines indicate single-trial trajectories. Subfigures indicate: (a) kinematic trajectories of cursor velocities for each reach direction; (b) neural trajectories in the 2-dimensional subspace identified by LDA; (c) neural trajectories in the 2-dimensional subspace identified via PCA; (d) kinematic trajectories of cursor positions for each reach direction; (e) reconstructed cursor positions from LDA trajectories; and, (f) reconstructed cursor positions from PCA trajectories. Neural trajectories in both the LDA (b) and PCA (c) subspaces only weakly correlate to true hand velocities (a). However, a linear control system reconstructs cursor positions with much higher correlations to observed kinematics. Reconstructed cursor positions from the LDA space (e) are more representative of true cursor positions (d) than those reconstructed from the PCA space (f).

**Supplementary Fig. 2:**
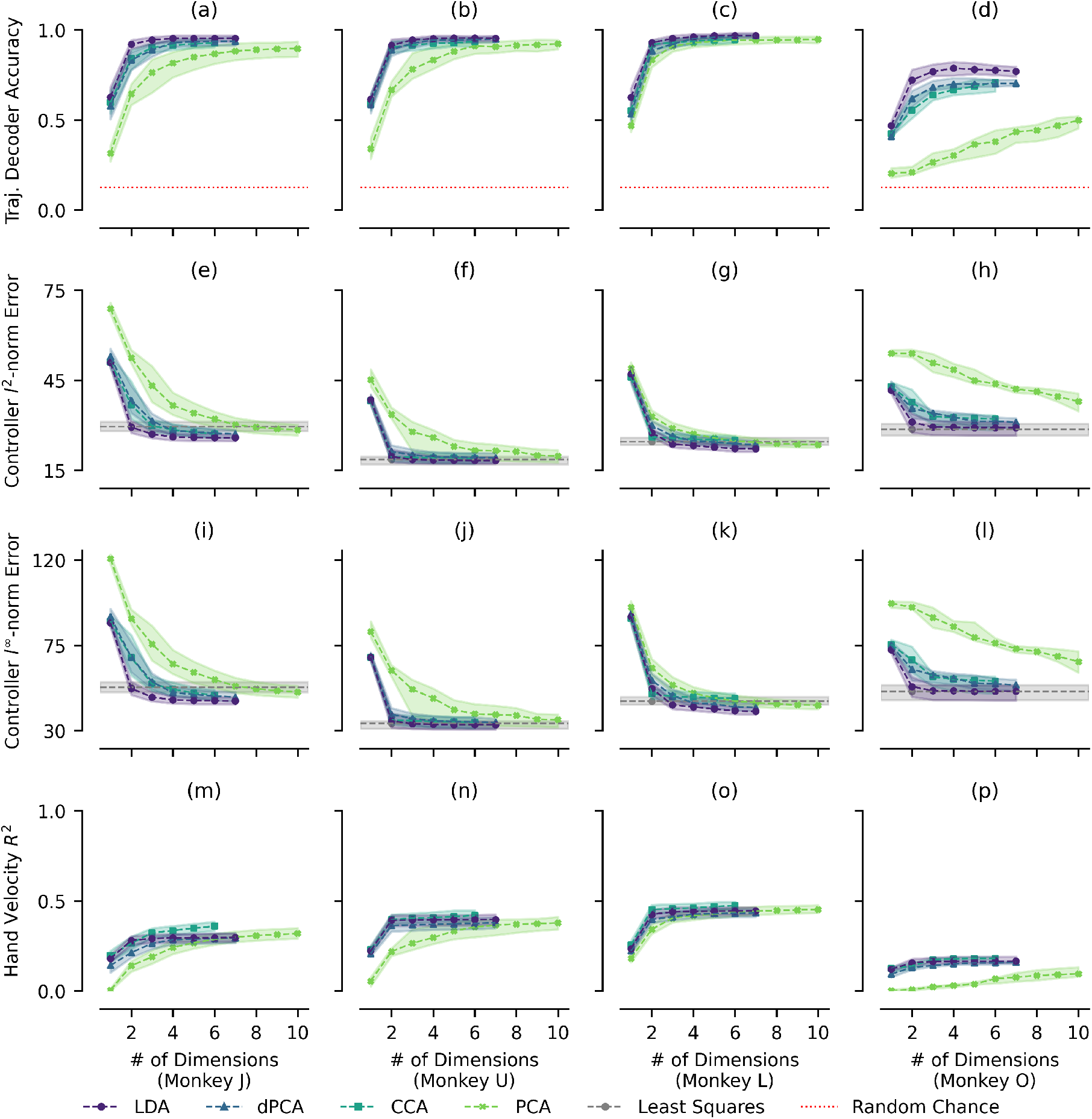
Alternative evaluation criteria for performance of dimensionality reduction methods at identifying behaviorally relevant latent variables vs. subspace dimensionality. Same as Figure 2, but here the rows indicate: decoding accuracy of a Gaussian Naive-Bayes decoder using the entire trajectory following the alternative methodology discussed in the Methods rather than individual samples (a, b, c, d), average *l*^2^-norm error of reconstructed vs. true cursor positions (e, f, g, h), average *l*^∞^-norm error of reconstructed vs. true cursor positions (i, j, k, l), and coefficient of determination with respect to hand velocities (m, n, o, p). Data was taken from all experimental sessions described in Supplementary Table 2.

**Supplementary Fig. 3:**
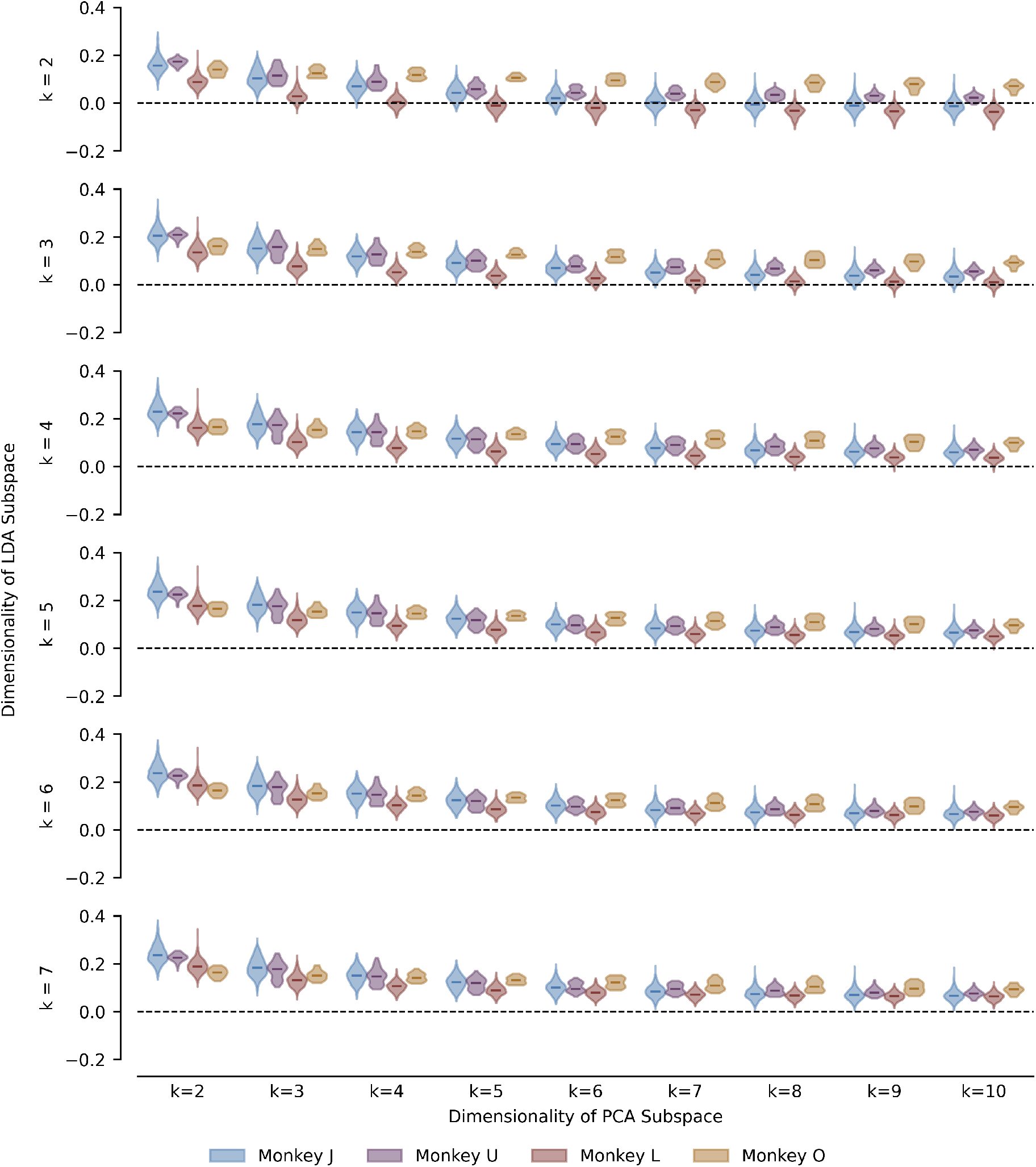
Net change in decoder accuracy when using LDA vs. PCA across various subspace dimensionalities. *Y*-axes represent the decoding accuracy of LDA minus the decoding accuracy of PCA. Black-dotted line indicates net 0 change in decoding accuracy. Rows and columns indicate subspace dimensionalities for LDA and PCA, respectively. For each row/column pair, four distributions are shown (one for each Monkey *J, U, L*, and *O*). Across all Monkeys, LDA results in better decoding accuracy with significantly fewer dimensions than PCA, although this difference approaches 0 as PCA dimensionality increases. Data is taken from the same experiments as Figure 2.

**Supplementary Fig. 4:**
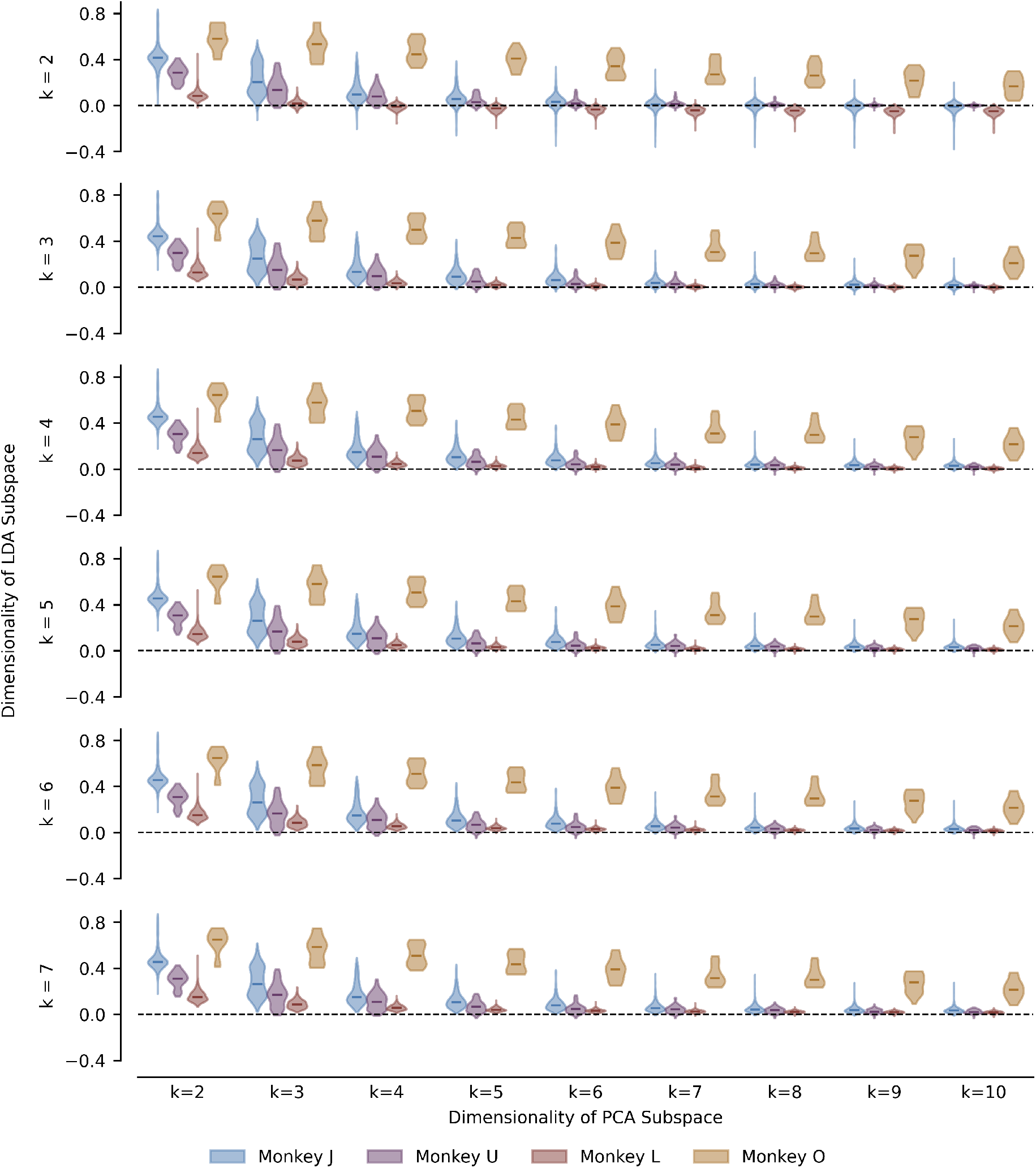
Net change in coefficient of determination between reconstructed and true kinematics when using LDA vs. PCA across various subspace dimensionalities. Same as Supplementary Figure 3, only *y*-axes indicate net change in cursor controller *R*^2^. Similar trends hold, where LDA with only 2-dimensions outperforms PCA with significantly more. Data is taken from the same experiments as Figure 2.

**Supplementary Fig. 5:**
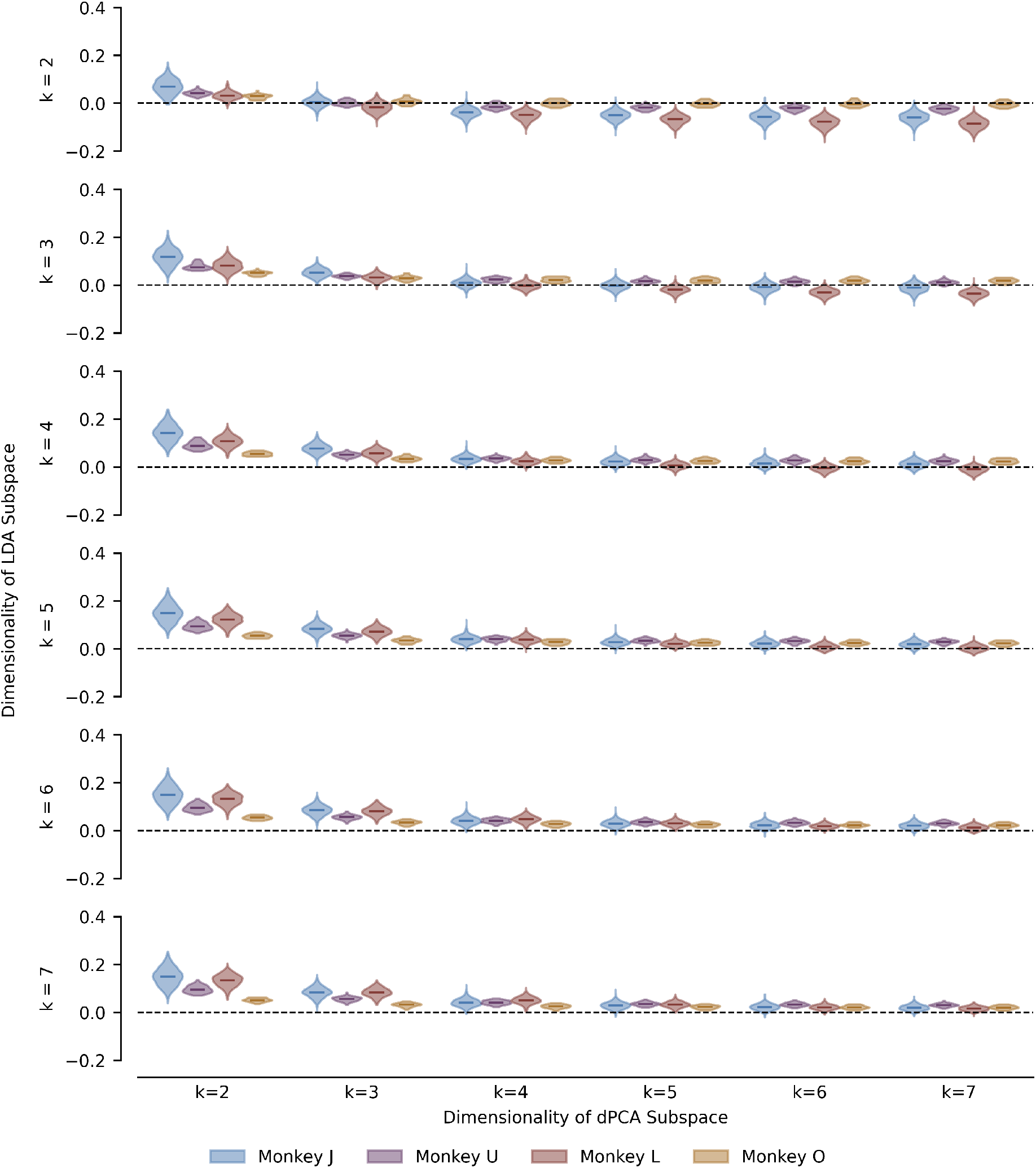
Net change in decoder accuracy when using LDA vs. dPCA across various subspace dimensionalities. Same as Supplementary Figure 3, only columns represent dimensionality of dPCA subspace, and *y*-axes indicate net change when compared to performance of dPCA. Same trends hold, although dPCA takes fewer dimensions to equalize performance with 2-dimensional LDA (Row 1). Data is taken from the same experiments as Figure 2.

**Supplementary Fig. 6:**
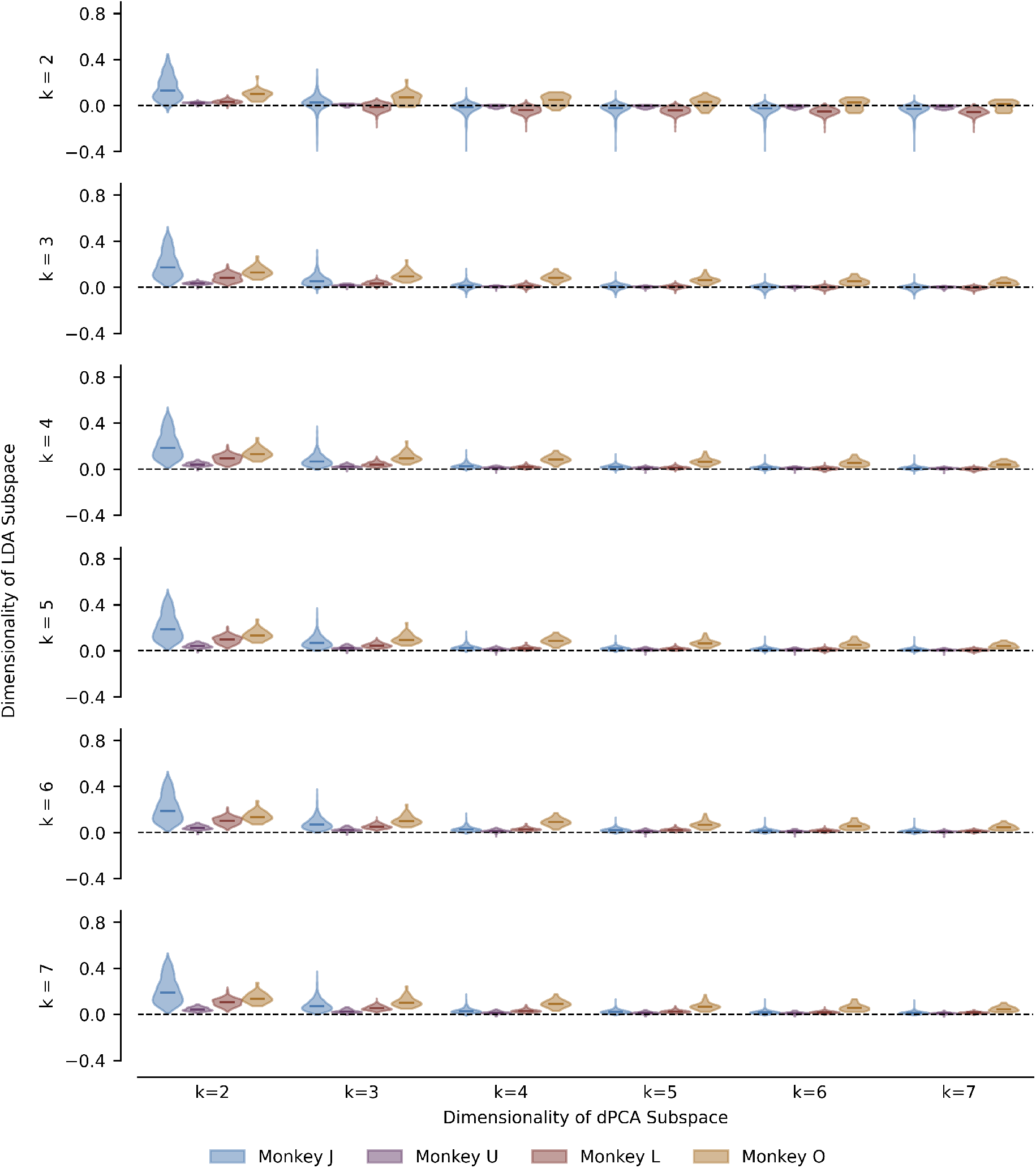
Net change in coefficient of determination between reconstructed and true kinematics when using LDA vs. dPCA across various subspace dimensionalities. Same as Supplementary Figure 5, only *y*-axes indicate net change in cursor controller *R*^2^. Same trends hold. Data is taken from the same experiments as Figure 2.

**Supplementary Fig. 7:**
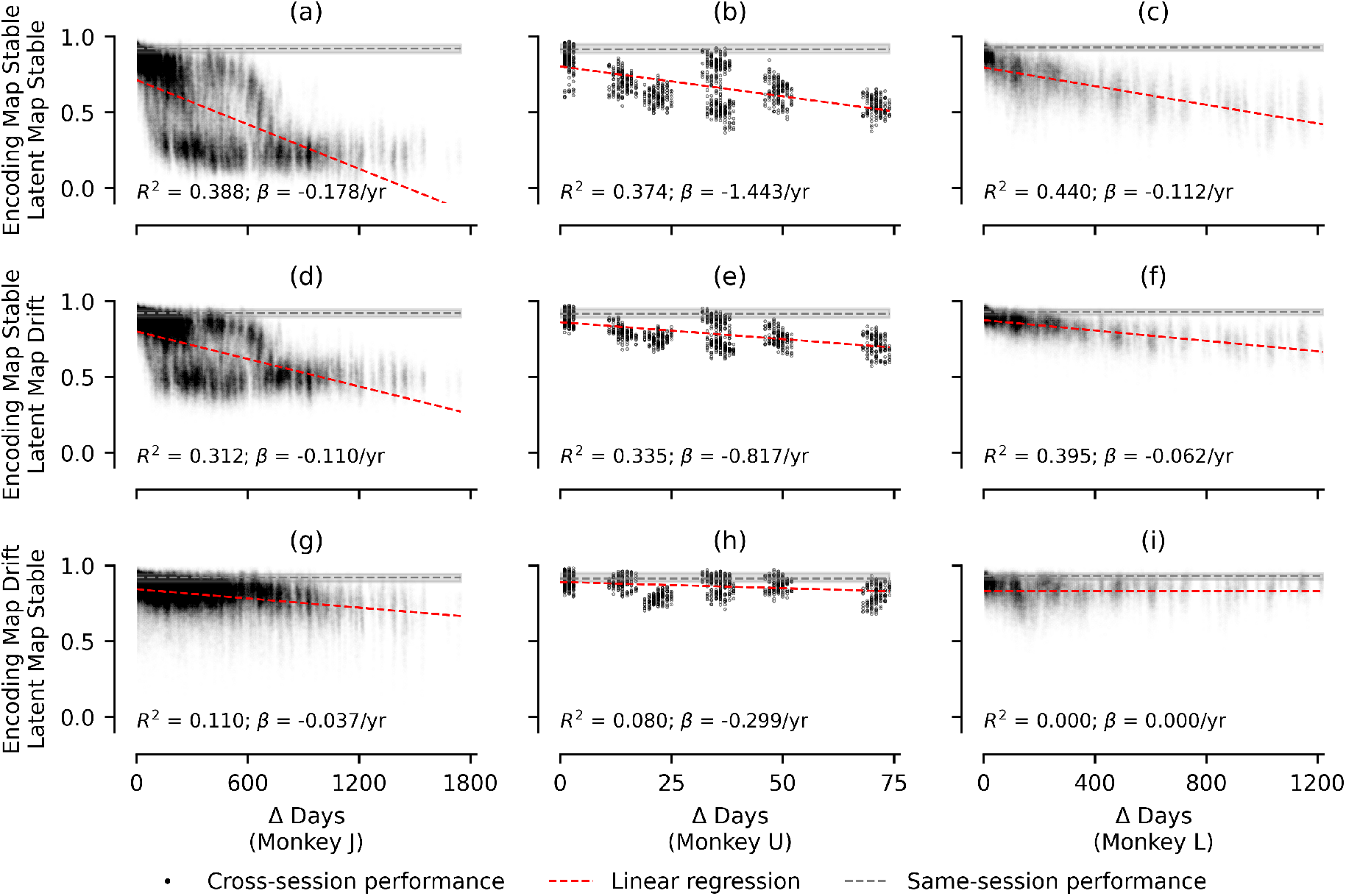
Cross-session trajectory decoding accuracy vs. number of days between sessions. Same as Figure 3, except *y*-axis of each subplot depicts the accuracy rate of classifying intended reach direction using entire latent trajectory on Day *j* using a Gaussian Naive-Bayes decoder trained on Day *i*.

**Supplementary Fig. 8:**
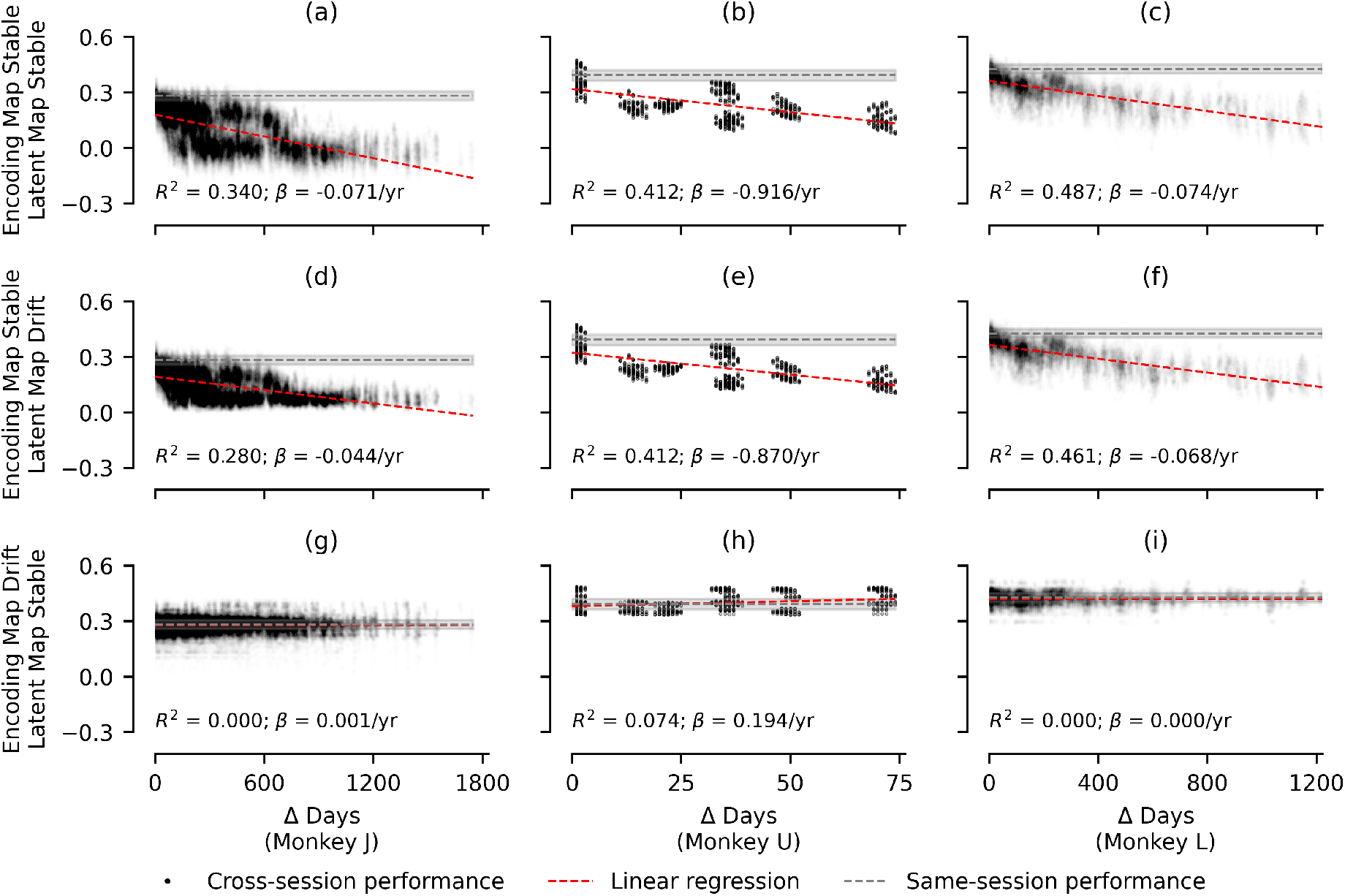
Cross-session hand velocity *R*^2^ vs. number of days between sessions. Same as Figure 3, except *y*-axis of each subplot depicts the coefficient of determination between latent state and instantaneous hand velocity on Day *j* using linear model trained on Day *i*.

**Supplementary Fig. 9:**
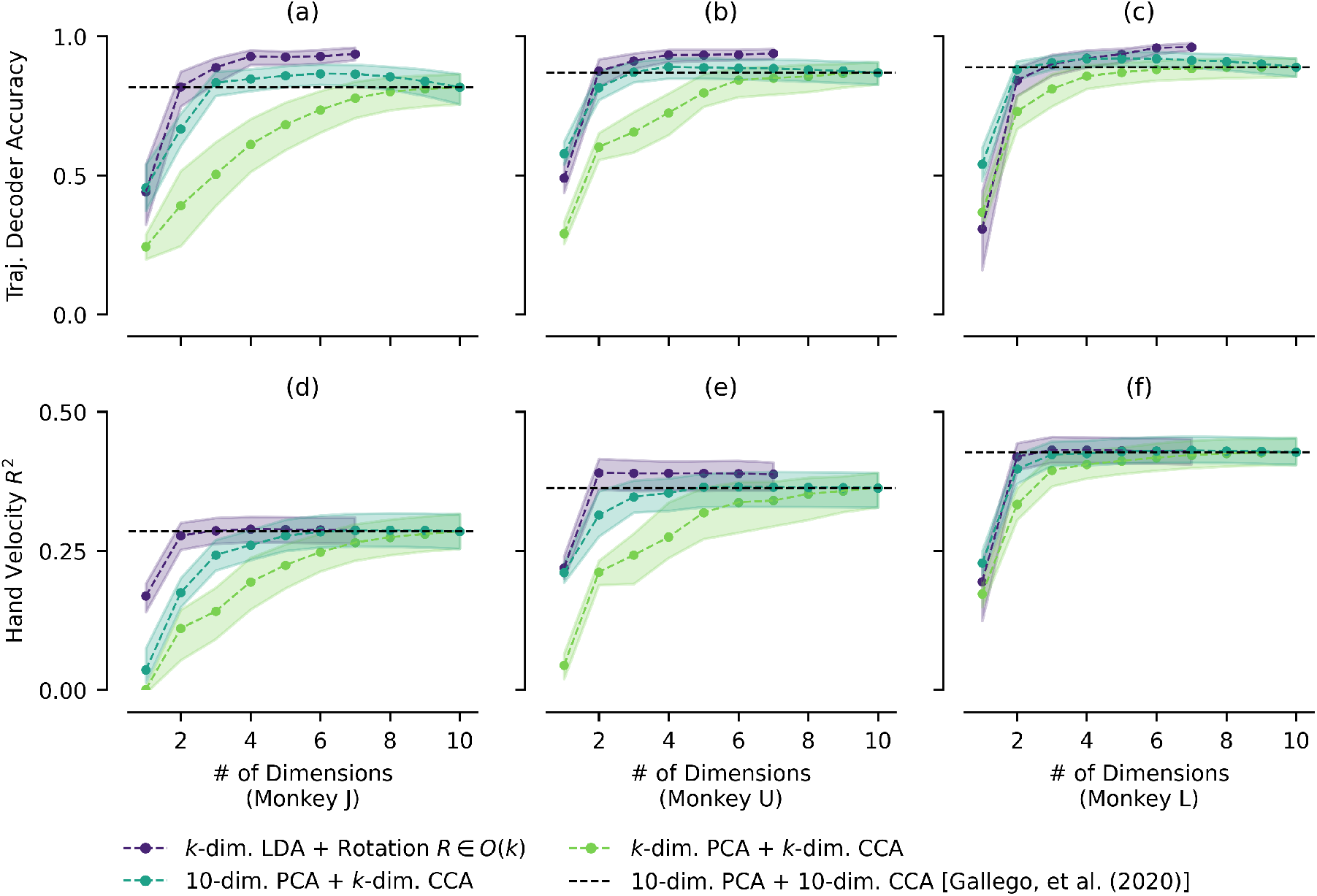
Alternative evaluation criteria for cross-session performance vs. subspace dimensionality using frameworks for stable long-term performance. Same as Figure 5, except *y*-axes reflect decoding accuracy using entire latent trajectory (a, b, c) and coefficient of determination with hand velocity (d, e, f). Similar to Figure 5, performance of the method proposed in this work with *k* = 2 matches that of the model proposed in Gallego, et al. (2020) ^39^ with *k* = 10.

**Supplementary Table 3:**
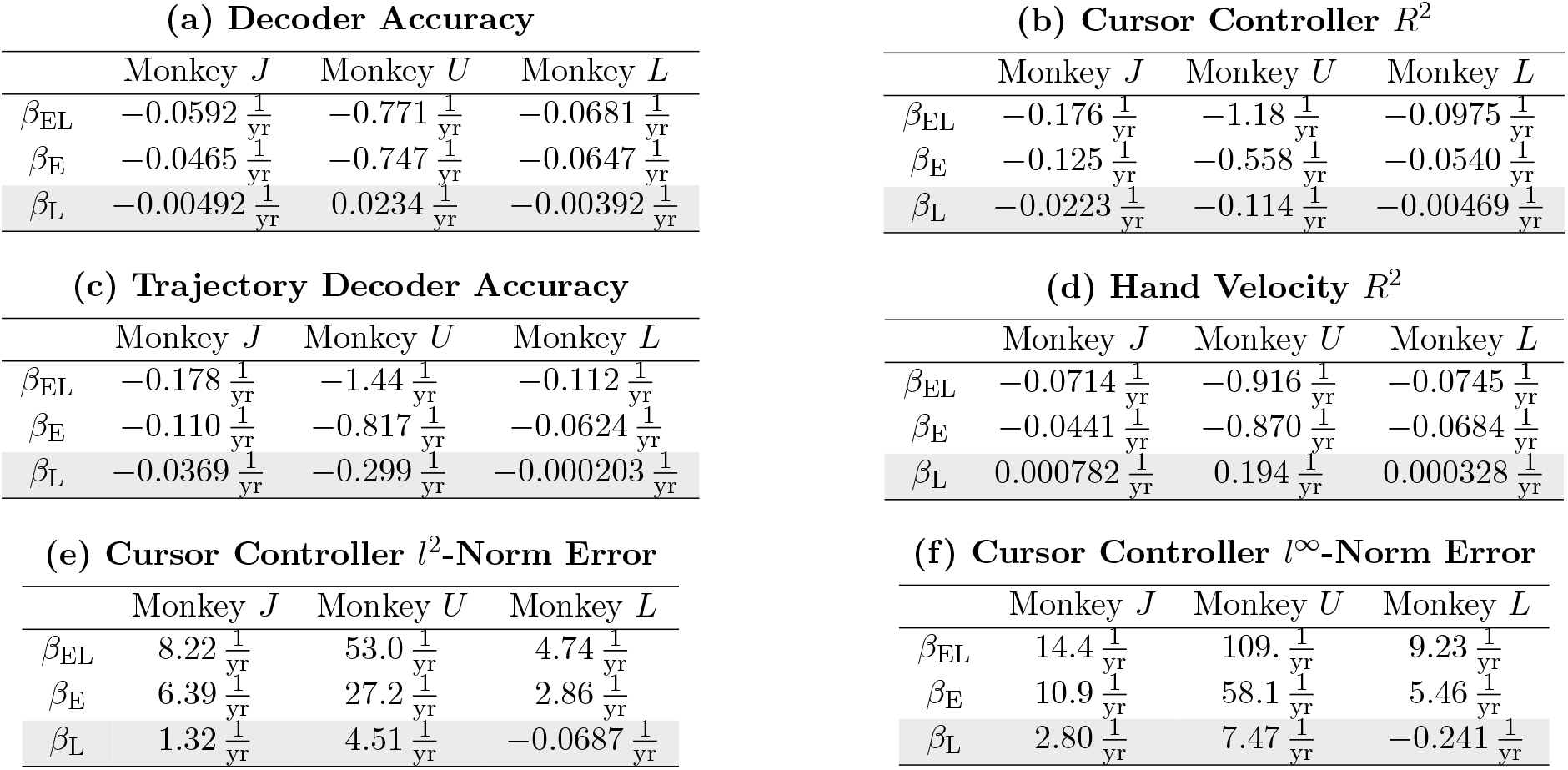
Model performance drift rates with 2-dimensional LDA subspace for various evaluation criteria under different hypotheses. Each subtable represents one of six evaluation criteria: (a) decoder accuracy; (b) cursor controller *R*^2^; (c) trajectory decoder accuracy; (d) hand velocity *R*^2^; (e) cursor controller *l*^2^-norm error; and (f) cursor controller *l*^∞^-norm error. Columns indicate performance drift rates for Monkeys *J, U*, and *L*, respectively. Rows indicate performance drift rates for models where both encoding and latent maps are held fixed (*β*_EL_), only encoding maps are held fixed while latent maps are retrained (*β*_E_), and only latent maps are held fixed while encoding maps are retrained and aligned (*β*_L_). All drift rates are calculated using linear regression on long-term performance of latent variables identified via 2-dimensional LDA. The model with drifting encoding maps and stable latent maps (grayed background) is the framework proposed in this work, which experiences drift in performance roughly one order of magnitude lower than the other proposed frameworks across all evaluation criteria.

**Supplementary Fig. 10:**
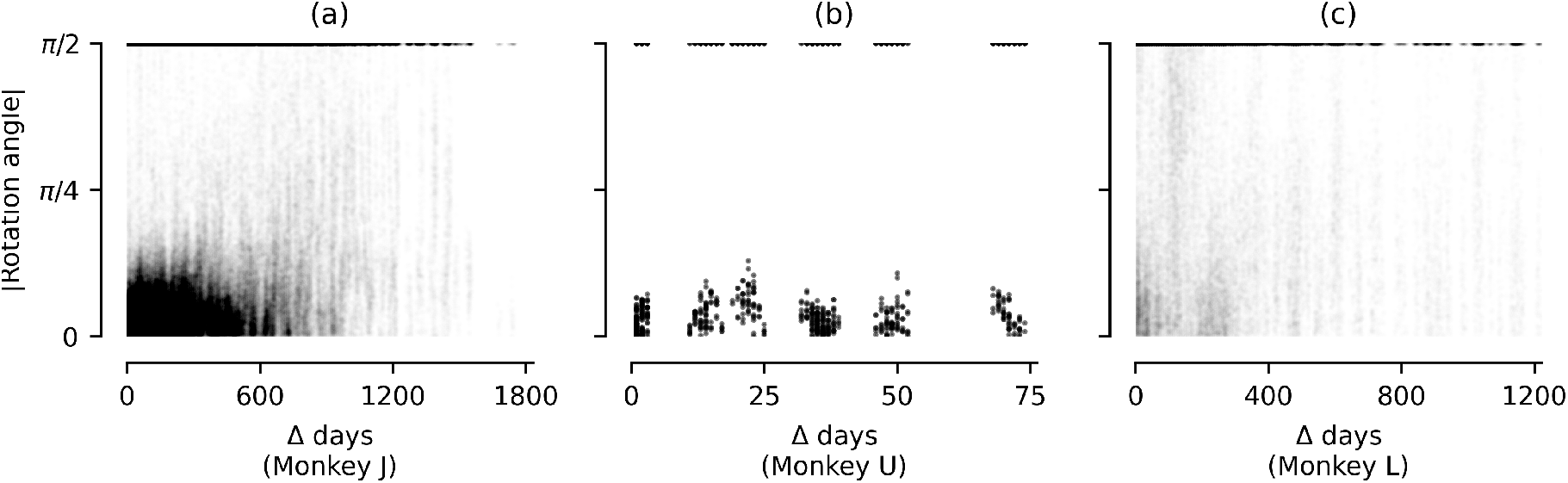
Magnitude of rotation angle of cross-session alignment rotation matrix *R* ∈ *O*(2) between *k* = 2-dimensional LDA subspaces vs. number of days between experimental sessions. Given a 2-by-2 rotation matrix *R*, angle magnitude was calculated using acos 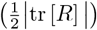. (a) Monkey *J*; (b) Monkey *U*; and, (c) Monkey *L*. Data was taken from same experimental sessions as in Figures 3 & 4.

**Supplementary Fig. 11:**
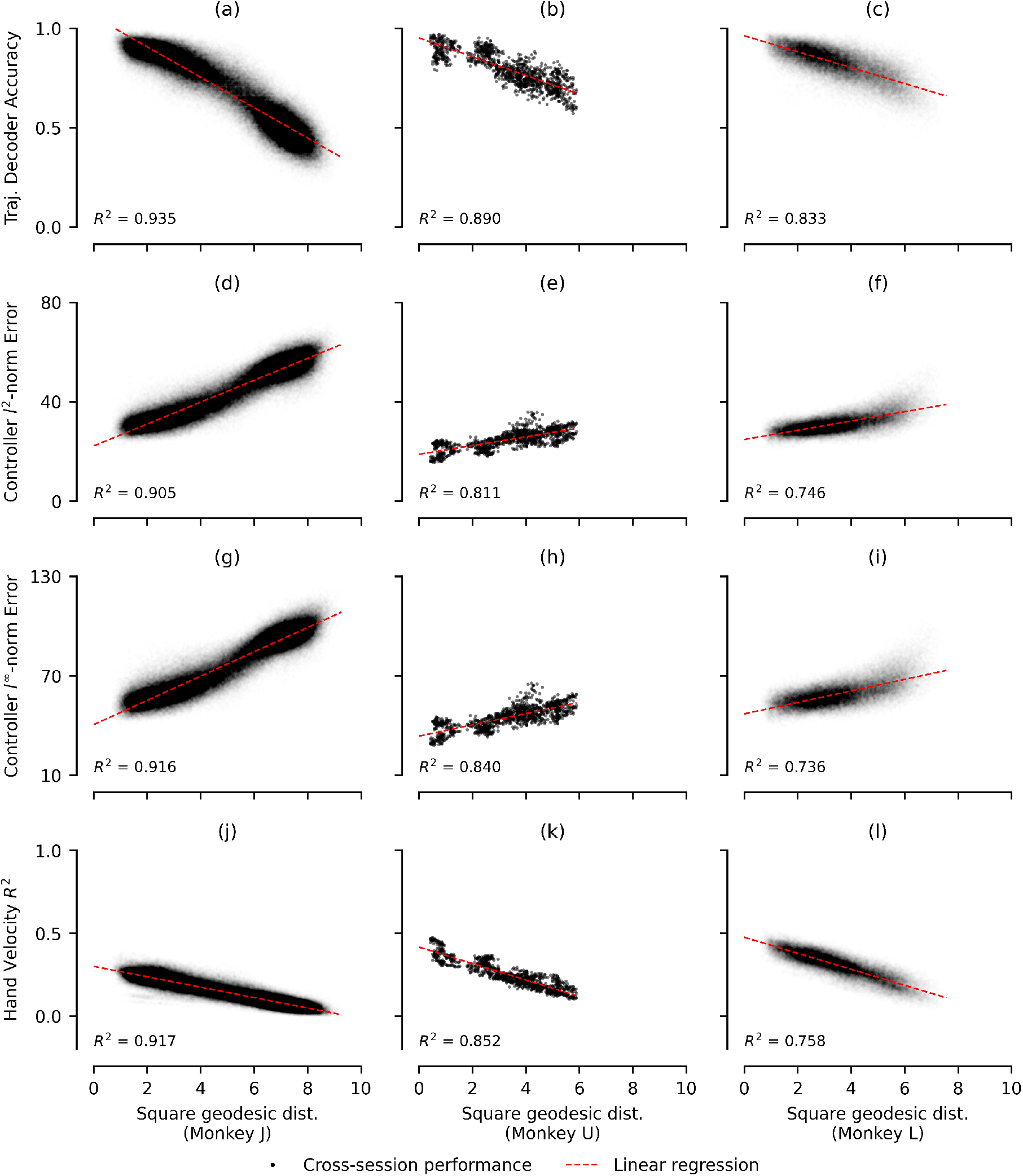
Cross-session performance using alternative evaluation metrics vs. geodesic distance between LDA subspaces between sessions. Same as Figure 6, with rows indicating accuracy of trajectory decoder (a, b, c), average *l*^2^-norm error of reconstructed vs. true cursor kinematics (d, e, f), average *l*^∞^ error (g, h, i), and coefficient of determination with respect to hand velocity (j, k, l). Data was taken from same experimental sessions as in Figure 3.

**Supplementary Fig. 12:**
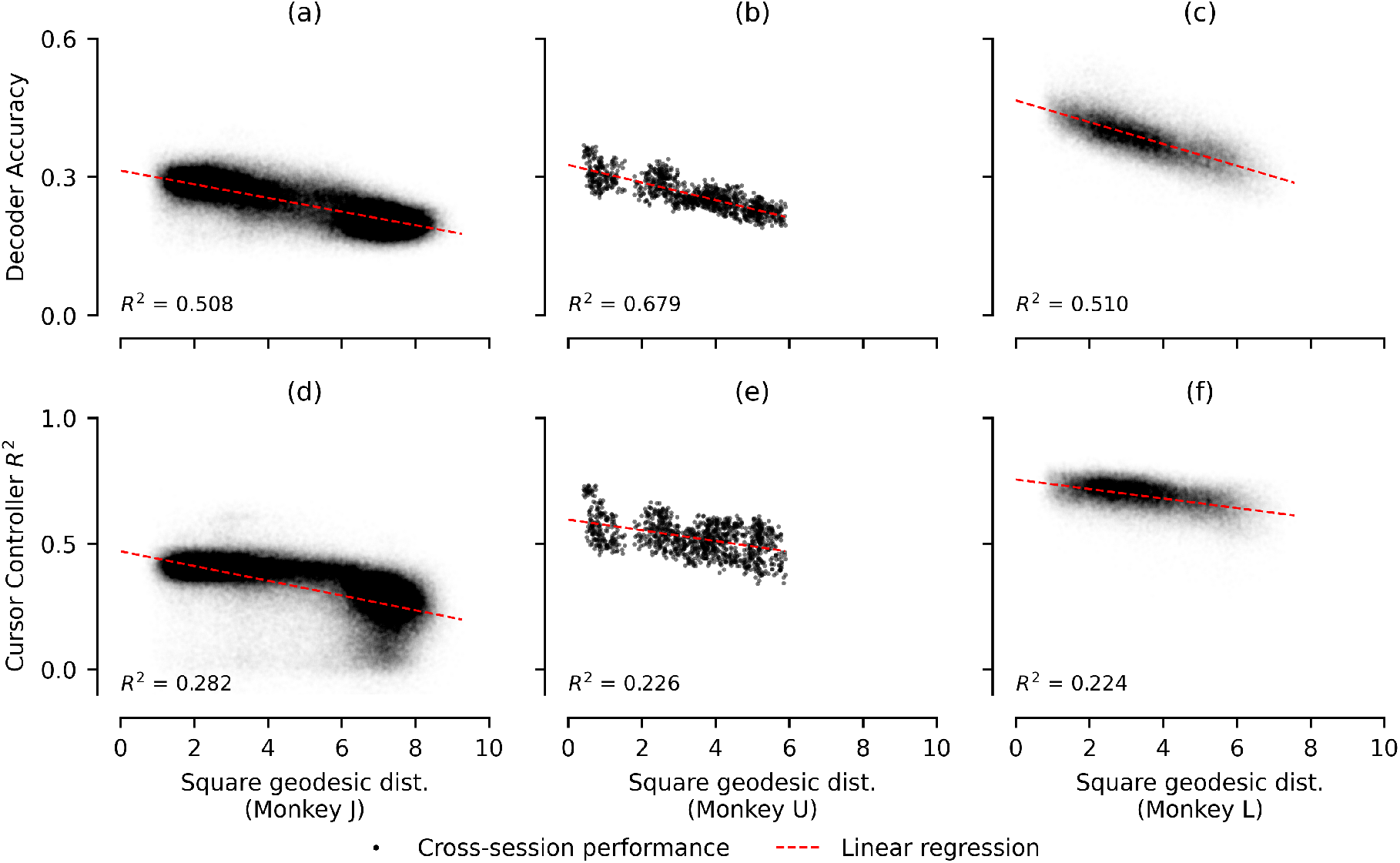
Cross-session performance vs. geodesic distance between 2-dimensional PCA subspaces between sessions. Same as Figure 6, only cross-session performance was measured using 2-dimensional PCA to identify the latent space, rather than 2-dimensional LDA. Geodesic distance was calculated between PCA subspaces. Note that compared to Figure 6, correlations are lower. This is most likely because 2-dimensional PCA does not capture the behaviorally relevant subspace.

**Supplementary Table 4:**
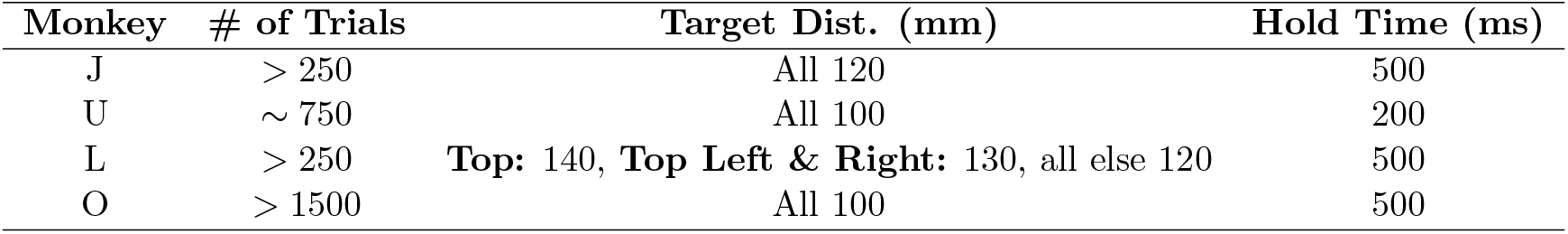
Summary of experimental parameters per monkey. Datasets from Monkeys *J* and *L* were filtered to only keep experimental sessions where *>* 250 trials were achieved in a single contiguous block of trials. Monkey *U* was performed 750 trials as part of the designed experimental paradigm. Monkey *O* was encouraged with a juice reward to perform as many trials as possible, with each session featuring between 1692 and 2327 total trials. Targets were arranged at radially at top, top left, left, bottom left, bottom, bottom right, right, and top right positions. For Monkey *L*, the top, top left, and top right targets were placed at greater distances from the center position than the remaining targets. For all other monkeys, targets were placed equidistant from the center point. Each monkey was expected to hold the cursor within a bounding box around the target position for a fixed amount of time for the trial to be considered successful.

| Monkey | # of Trials | Target Dist. (mm) | Hold Time (ms) |
| --- | --- | --- | --- |
| J | > 250 | All 120 | 500 |
| U | ~ 750 | All 100 | 200 |
| L | > 250 | **Top: 140, Top Left & Right: 130, all else 120** | 500 |
| O | > 1500 | All 100 | 500 |

**Supplementary Fig. 13:**
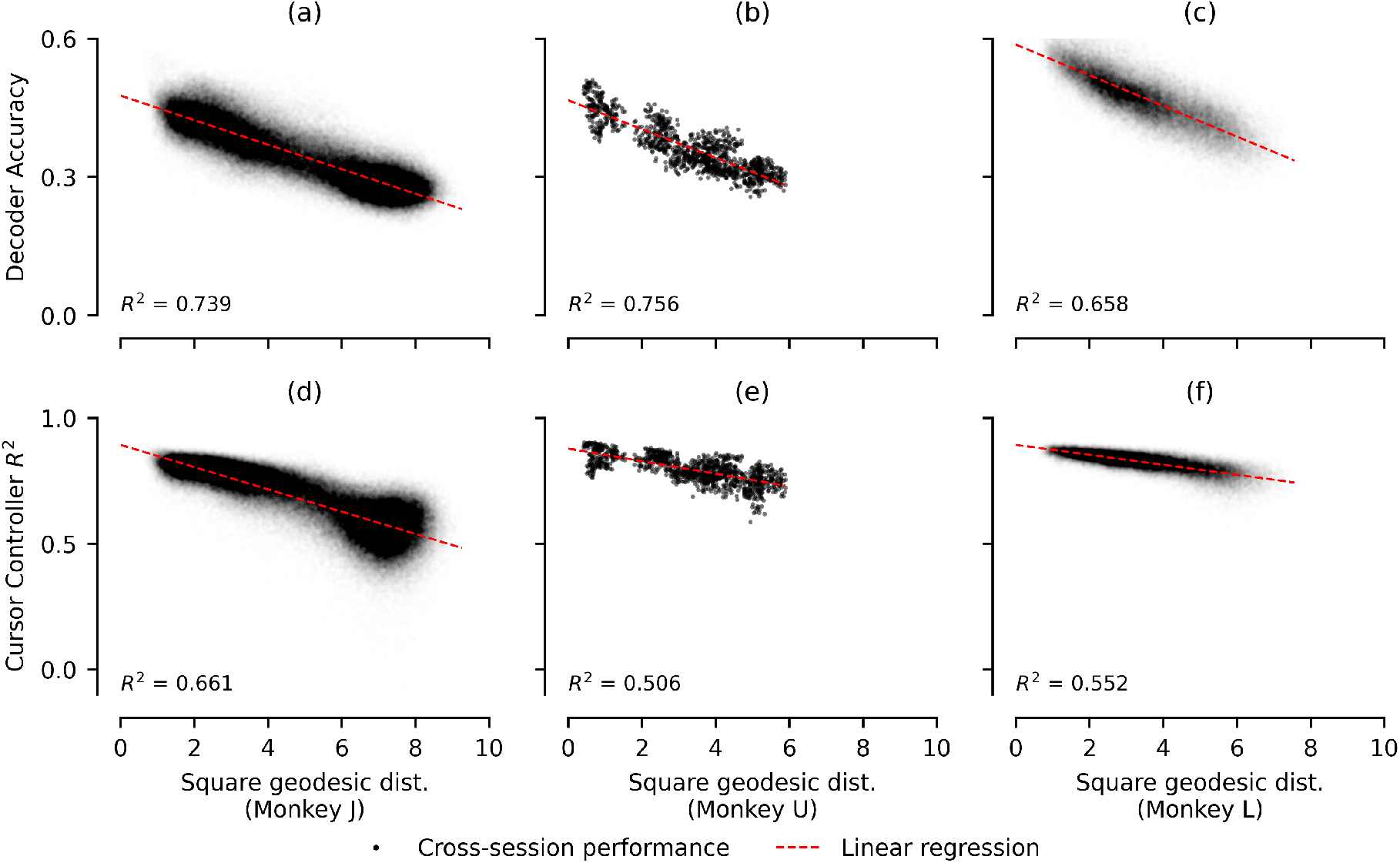
Cross-session performance vs. geodesic distance between 10-dimensional PCA subspaces between sessions. Same as Figure 6, only cross-session performance was measured using 10-dimensional PCA to identify the latent space, rather than 2-dimensional LDA. Geodesic distance was calculated between PCA subspaces. Note that compared to Figure 6, correlations are relatively similar (only slightly lower.) This is most likely because 10-dimensional PCA does span the behaviorally relevant subspace, but also captures additional dimensions.

**Supplementary Fig. 14:**
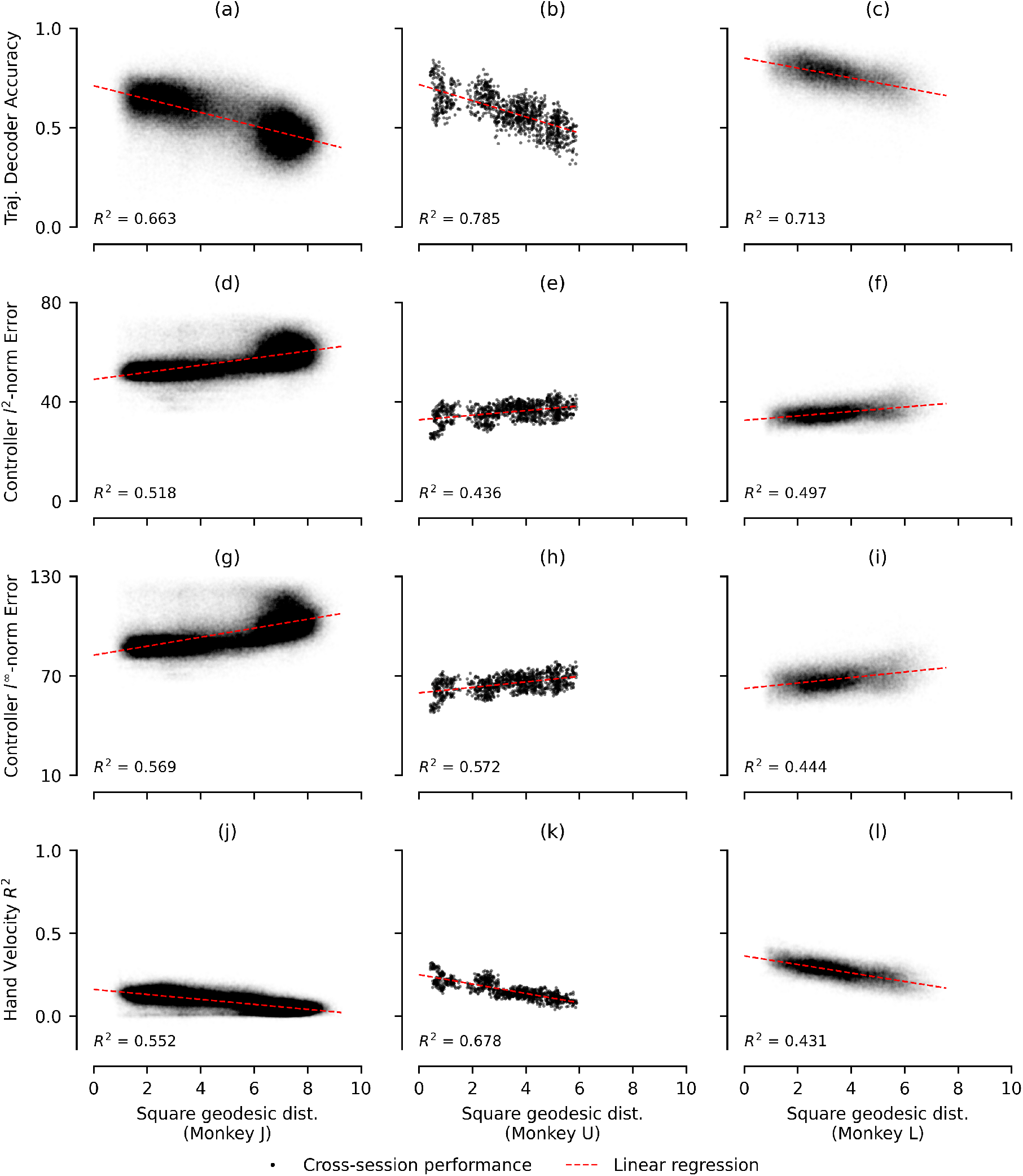
Cross-session performance using alternative evaluation metrics vs. geodesic distance between 2-dimensional PCA subspaces between sessions. Same as Supplementary Figure 11, only cross-session performance was measured using 2-dimensional PCA to identify the latent space, rather than 2-dimensional LDA. Geodesic distance was calculated between PCA subspaces. Again, correlations are much lower.

**Supplementary Fig. 15:**
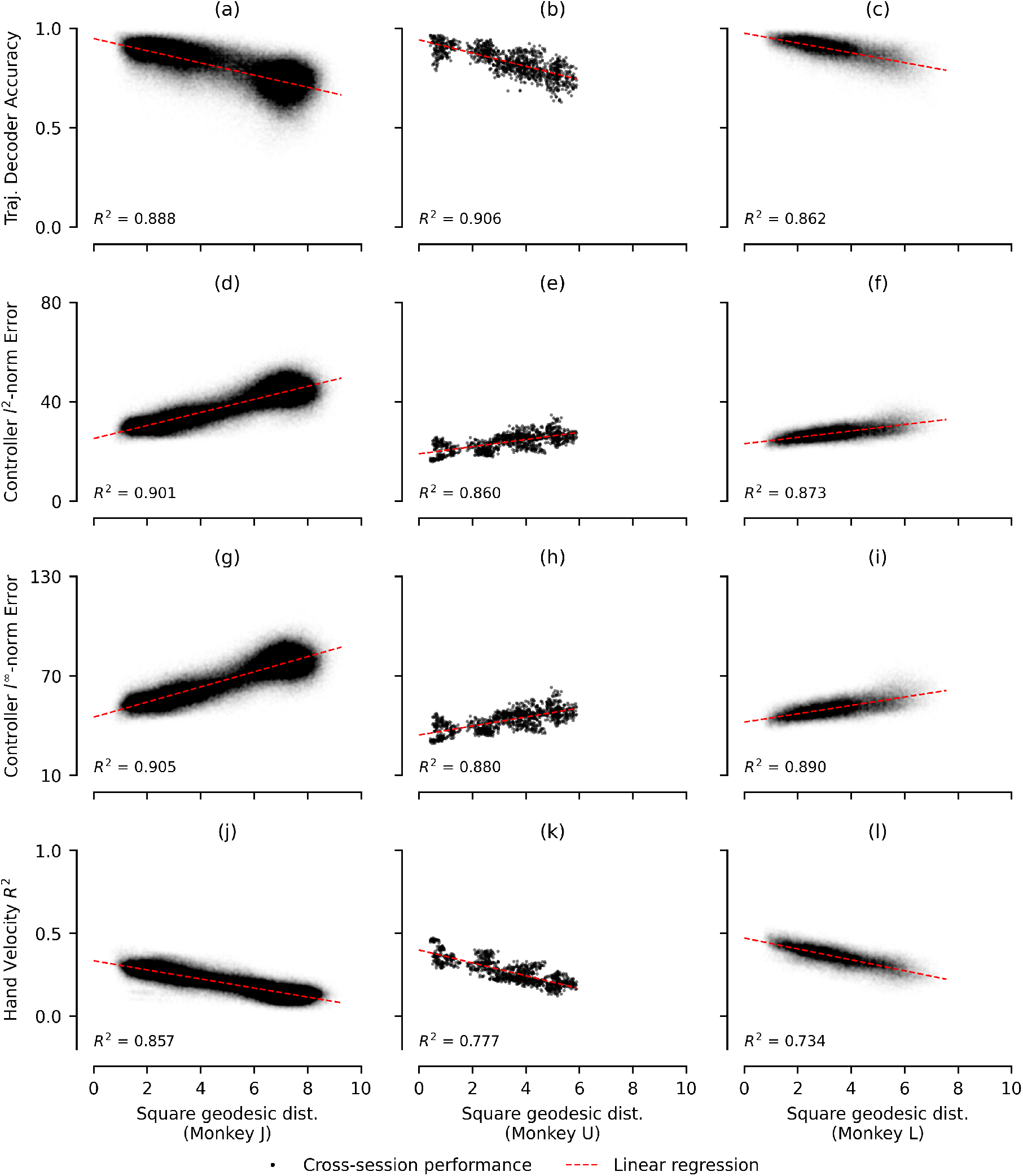
Cross-session performance using alternative evaluation metrics vs. geodesic distance between 10-dimensional PCA subspaces between sessions. Same as Supplementary Figure 11, only cross-session performance was measured using 10-dimensional PCA to identify the latent space, rather than 2-dimensional LDA. Geodesic distance was calculated between PCA subspaces. Again, correlations are similar.

**Supplementary Fig. 16:**
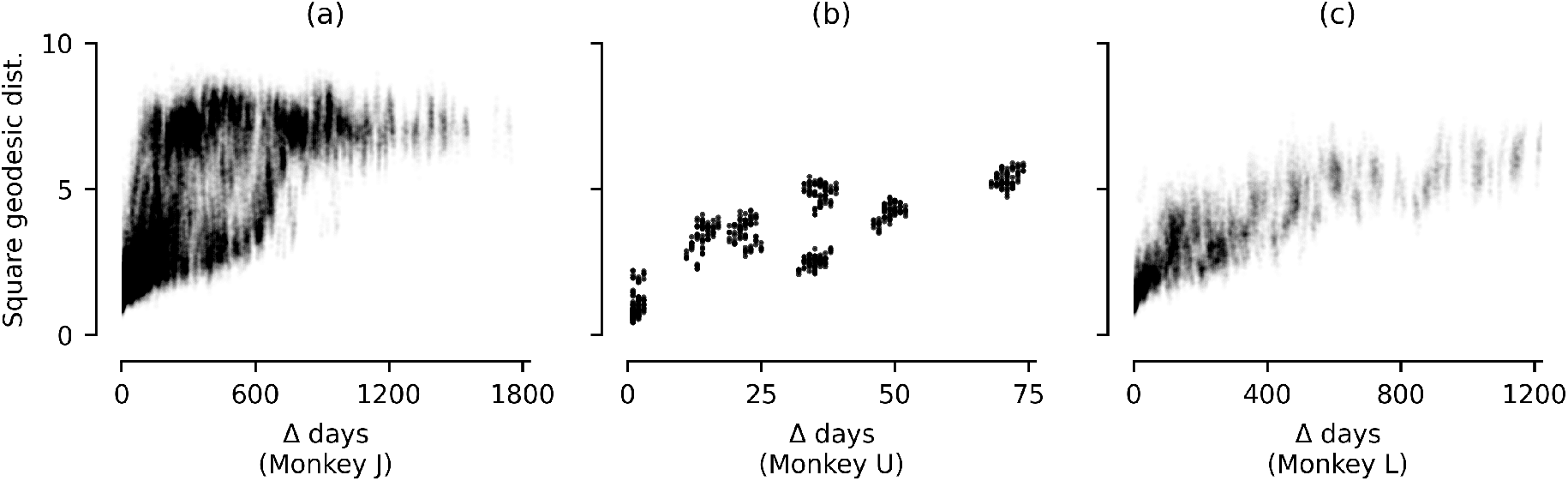
Geodesic distances between 2-dimensional encoding maps (identified via LDA) vs. number of days between sessions. *Y*-axis indicates square geodesic distance between subspaces identified on Days *i* ≠ *j* and *x*-axis indicates absolute difference in time |*i*− *j*|. Subplots depict results for Monkeys *J* (a), *U* (b), and *L* (c), respectively. Col. 3: Monkey *L*. Data was taken from same experimental sessions as in Figure 3.

**Supplementary Fig. 17:**
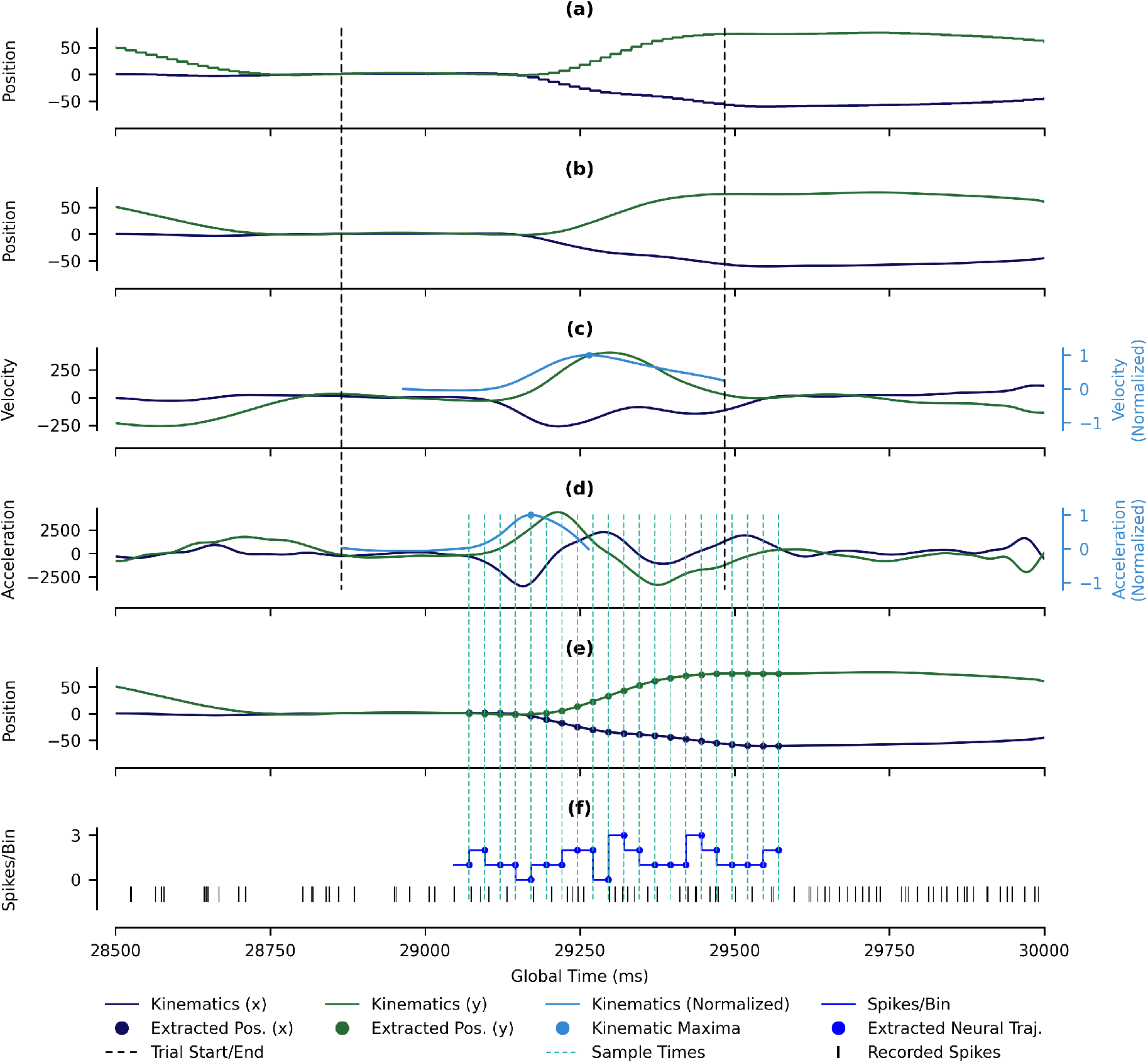
Depiction of the preprocessing and alignment algorithm used for a sample reaching trial for Monkey *U*. (a) Raw cursor position is updated at a sample rate of 60 Hz (with *x* in navy blue, *y* in green). Trial start (time when target is shown) and end (time when target is reached) indicated in dotted black. (b) Smoothed cursor position data (with *x* in navy blue, *y* in green). (c) Smoothed cursor velocity data (with *x* in navy blue, *y* in green, and “normalized” in solid light blue). (d) Smoothed cursor acceleration data (with *x* in blue, *y* in green, and “normalized” in solid light blue). Trial is “centered” according to point of peak normalized acceleration. (e) Kinematics extracted by sampling smoothed position data every 25 ms (dotted sky blue lines). (f) Kinematics are paired with samples of neural data (blue dots) by counting the number of spikes (solid black) observed in the 25 ms bin (blue horizontal lines) immediately preceding each kinematic sample (green and navy blue dots).

